# Living with urban fear: vervet monkey response to an evolutionarily new predator

**DOI:** 10.1101/2025.10.04.679731

**Authors:** Benjamin Robira, Natacha Bande, Stéphanie Mercier, Andri Manser, Charlotte Vanderlocht, Sofia Forss

**Affiliations:** Department of Evolutionary Biology and Environmental Studies, University of Zurich, Switzerland; Faculty of Life Science, University of Strasbourg, France; School of Life Sciences, University of KwaZulu-Natal, South Africa; Department of Biological, Geological and Environmental Sciences, University of Bologna, Italy

**Keywords:** domestic dogs, memory, prey naïveté, predator recognition, risk labels, stimulus discrimination

## Abstract

Humans have facilitated contacts between prey and predator species that have originally not co-evolved, reshuffling the prey-predator arms race. How do prey cope with an evolutionarily new predation risk? We tracked three vervet monkey troops in a South-African semi-urban habitat for 14 months to study their response to domestic dogs. We show that monkeys responded to dogs with a two-pronged behaviour: they emitted alarm calls, and became more vigilant and displayed aggressive behaviours towards the dogs. While their movement highlighted risk-prone behaviour, they appeared to have mapped and planned for risk, as they reacted more strongly when risk was unexpected. The response intensity was further modulated by risk labels typically encountered in their natural environment, but not by labels uniquely associated with dogs. This highlights that vervet monkeys responded with ingrained behaviour to this evolutionarily new threat, anticipating risk based on long-term spatial memory, but failed to integrate evolutionarily new information.

**Teaser:** Vervet monkeys fear domestic dogs but fail to adjust their antipredator response by discriminating evolutionarily new risk labels.

## Introduction

Prey and predators are engaged in an evolutionary arms race: an arsenal of defensive traits has been favoured in prey in response to traits shaping predators’ hunting strategies, resulting in increased selection pressure on further (anti)predator traits, and so on, following “Red Queen dynamics” (*1*). In this ongoing race, prey defensive responses are behaviourally and functionally organised along the predation sequence (*2*) - avoidance (in space and time, through adjustment of space use and activity patterns), detection (e.g., vigilance and environment monitoring), and pursuit- deterrence (e.g., alarm calls, postural displays, or agonistic behaviours) - all of which rely on cognitive processing of risk information (*3*). In existing empirical research, however, responses are often grouped by data domain: vocal responses (based on acoustic monitoring), directly observable interactional behaviours (i.e., proximal defensive behaviours directed at the predator, and identified based on field observations, such as alertness, fleeing, or agonistic displays), and movement-based responses (based on remote-sensing tracking). An integrated analysis of vocal, interactional and movement behaviours offers insights into prey defensive strategy and their adaptation to (new) predation risk.

### The vocal, interactional, and movement response of prey

Vocally, prey may advertise the predator’s presence. In gregarious species, these calls may alert conspecifics, possibly inducing increased alertness, flight or a defensive tactic (*4*), or they may signal to the predator itself that it has been spotted (*5*, *6*) and potentially serve an intimidating function (*7*). For communicative purposes, not all vocalisations are alike, and each may encode specific information. For example, vervet monkeys (*Chlorocebus pygerythrus*) have alarm calls that are produced towards specific predator types (*8*), inducing predator-specific behavioural responses [climbing a tree for leopards, standing up bipedally for snakes or running to bushes for raptors (*9*)].

With regards to direct interaction with predators, prey may also become more vigilant and search for signs of predators when the timing (*10*) or location (*11*) is favourable for predator attacks. However, interactional responses are not limited to proactive behaviours; prey may also respond to a detected predator (Crane et al., 2024) with reactive agonistic behaviour. For instance, muskoxen (*Ovibos moschatus*) often retaliate against wolves (*Canis lupus*) by collectively forming an arc of protection surrounding the weakest individuals and charging the canids (*12*).

From a movement perspective, prey may proactively avoid high-risk areas (*13*), reactively flee when a predator is spotted (*14*) or, conversely, freeze (*15*) and possibly hide, or delay returning to previous encounter sites (*16*).

In any case, these responses can carry costs: vocalisations may attract distant predators (*17*), increased vigilance may reduce foraging efficiency and avoiding certain areas may mean forgoing valuable resources (*18*). To minimise the costs, prey must therefore calibrate their responses according to perceived risks.

### Prey’s risk perception abilities

Not all situations pose the same level of danger, and prey typically discriminate between low- and high-risk scenarios to adapt their responses efficiently (*19*). For example, when predators are spotted at a reasonable distance, prey may just increase vigilance. When, however, the predator crosses a certain distance threshold (*20*), prey may engage in fleeing or fighting. This distance may be shaped by factors affecting the degree of exposure that the prey may perceive (e.g., proximity of refuges, own elusiveness, (*21*)).

Risk, indeed, is understood as a product of both hazard and exposure. Prey continuously process external stimuli to assess the likelihood of predator presence, and how exposed they would be in case of danger (*22*). Both the evaluation and the resulting behaviour may rely on either innate or learned processing of labels (also referred to as cues, markers or features) of risk (*23*). For example, cuttlefish (*Sepia officinalis*) embryos can innately react to chemical and visual cues indicating the presence of predators, reducing their ventilation rate and therefore making them less visible. However, through associative rules involving an extrinsic risk signal (e.g., conspecific ink emission), they can also learn to recognise new dangers and adjust their defensive behaviour accordingly (*24*). In both cases, adaptive responses to predation involve a high degree of (innate) familiarity with (cues of) predators and their environment.

### An urbanised landscape of fear

Humans are increasingly transforming the world, hence creating evolutionary unfamiliar conditions due to climatic variations (*25*), habitat structure changes (*26*) and changes in species distribution (*27*). The alteration of species distribution may occur in ‘natural’ habitats, either due to voluntary or involuntary dispersal, but also in already human-altered habitats where domestic “exotic” animals are hosted and may thus cohabit with wild urban species. As a result, wild animals may encounter entirely novel threats, including potential predators they have never faced before, raising questions about the consequences of their naïveté (*28*), and their ability to assess and respond to new risk labels (*23*).

Facing novel predators in an urban environment may be challenging, as prey must integrate new labels of risk relating to both the predator itself, and the altered context of predation associated with anthropisation. For example, while domestic dogs may pose a threat to small- and medium-sized animals (*29*), not all dogs are equally dangerous. Despite belonging to the same species and showing broadly similar behaviours, domestic dogs indeed display far greater phenotypic diversity than natural predators, ranging from relatively harmless small dogs to more threatening hunting breeds (*30*). Moreover domestic dogs can be overweight or elderly, conditions rarely encountered in the wild, that reduce their motor and sensory abilities (as observed in captive species (*31*)), making them poor hunters.

Urban environments may also influence prey exposure. Despite the disturbances humans may cause (*32*, *33*), or the direct lethal risk they pose (*34*), some species benefit from human proximity, using it as a shield against predators (*35*). Urban infrastructure, houses, fences, and roads also alter movement dynamics, creating both barriers and shelters, or escape routes for prey (*36*). Furthermore, urban prey species must also incorporate human artefacts into their risk assessment. For instance, although dogs may roam freely, they may also be restrained by leashes or confined to fenced yards, reducing their threat level. However, prey may not necessarily know how to interpret these restrictions as reliable safety signals. Interestingly, some urban species, like cockatoos (*Cacatua galerita*), have been observed manipulating human artefacts (*37*), suggesting an ability to grasp their function and implications, raising the question of how well prey can adapt, or have adapted, to live with this urban fear.

### Study aims

In this study, we investigated how vervet monkeys living in Simbithi Eco-Estate (hereafter Simbithi), in Ballito, KwaZulu-Natal, South Africa, respond to the urban threat of domestic dogs. Vervet monkeys are naturally predated by pythons, raptors and ambush predators such as leopards (*9*). In Simbithi, although raptors, and, to a lesser extent, pythons, remain a natural threat, monkeys here inhabit an urbanised area where leopards are absent, but where human residents own domestic dogs, known to threaten many vertebrate species (*38*). These dogs may roam freely in gardens or be walked, with or (against the estate’s guidelines) without a leash, throughout the estate. Aligning with observations of several monkey species being attacked and even killed by domestic dogs in other anthropised areas of the world (*39–43*), several reports of dog attacks on vervet monkeys in our study area have indeed been made to the environmental team of Simbithi (Simbithi Eco-Estate Administration, unpublished data) in line with reports of vervet monkeys being injured or even killed by dogs in nearby areas (*44*). As Simbithi is only a few decades old, the habitat provides an experiment-like setting where vervet monkeys are still evolutionarily “naïve” to the local conditions. It therefore raises the question of how vervet monkeys cope with this new predator. Specifically, do vervet monkeys display antipredator responses toward this “evolutionarily novel” threat, and are they capable of discriminating between the various (novel) risk labels? To address these questions, we studied the monkeys’ behaviour during 755 encounters with domestic dogs, focusing on three aspects: vocal responses, interactional behaviours, and movement patterns, to assess whether they exhibited alarm calls, increased vigilance, and/or avoidance. Furthermore, we examined which dogs’ characteristics and micro-habitat features influenced the monkeys’ fear responses. If vervet monkeys have assimilated domestic dogs as a novel threat, then labels previously irrelevant in natural predation (e.g., number of predators - as their terrestrial predators are usually solitary, lowered physical health condition - since their natural predators are generally fit, and barking - unlike their silent natural predators) should elicit similar responses to familiar indicators of danger such as body size. The same logic applies to the micro-habitat, where new labels of exposure (e.g., human artefacts that constrain dog mobility) are added to archetype labels such as distance to the predator (horizontally or vertically).

## Methods

### Study site and species

The study was conducted within the Simbithi Eco-Estate, a 430-hectare gated residential eco-estate near Durban, in the province of KwaZulu-Natal, South Africa. Simbithi includes housing complexes and community centres dissected by roads and remnant forest patches, as well as artificial green areas. This forms a heterogeneous semi-urban mosaic landscape which hosts both wild and domestic animals.

We studied three vervet monkey troops that were habituated to human observers when the study was conducted from February 2024 to April 2025 (locally named *Acacia*, *Savanna*, and *Pink*, hereafter referred to as Troops 1, 2 and 3, respectively; Figure 1). Vervet monkeys from this eco-estate exploit both natural and anthropised areas and thus are in close contact with humans throughout the day. This raises human-wildlife conflicts (*45*) and exposes them to “new” domestic animals.

**Figure 1:**
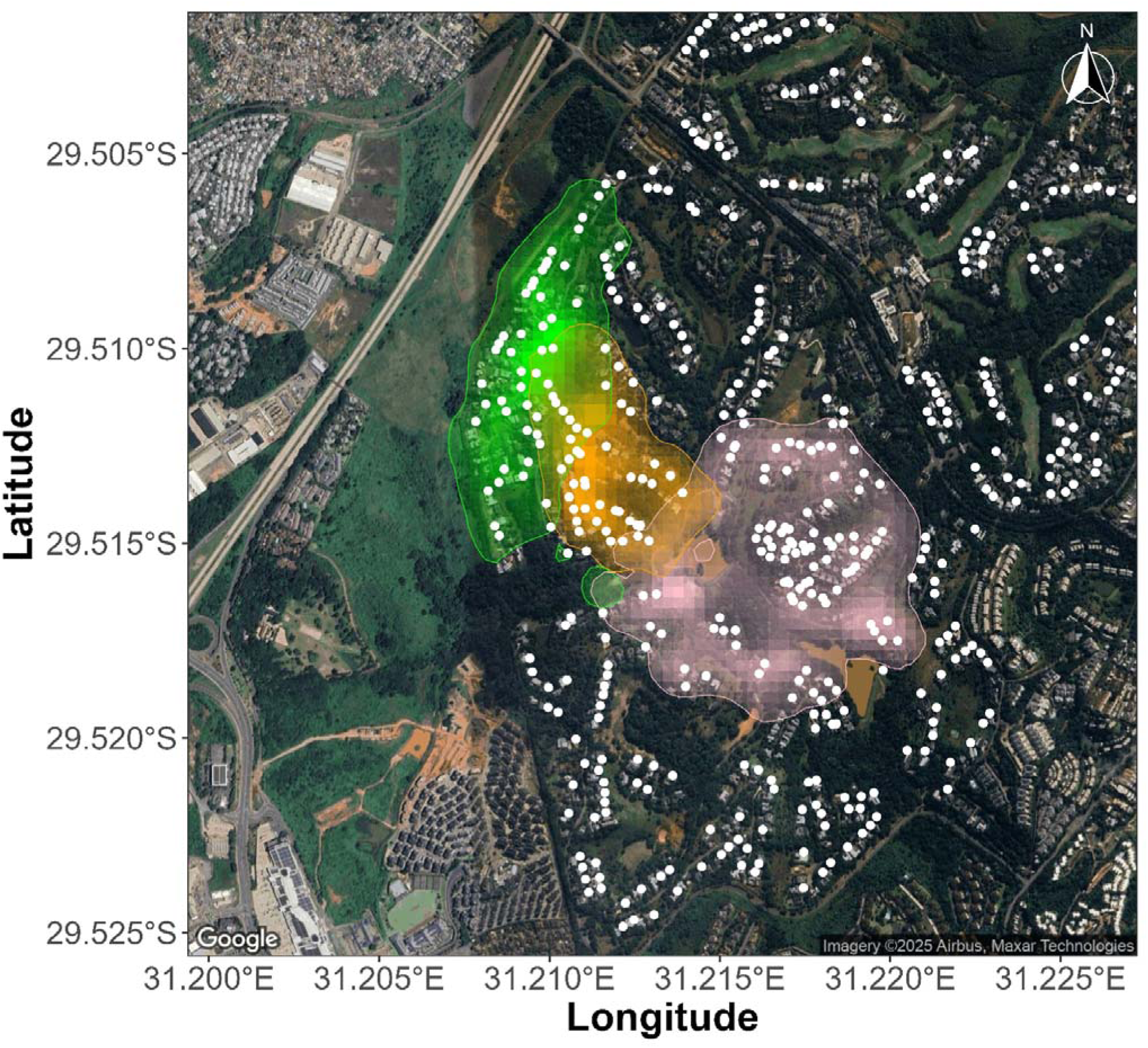
Study monkey troops’ ranges in Simbithi Eco-Estate. The range of each troop is delineated by the plain line. Monkey troops’ ranges corresponded to 95% of their spatial utilisation distributions, which were estimated using Brownian-based movement kernels (*46*) with the “BRB” function of the *adehabitatHR* package (*47*). The intensity of colouration reflects the intensity of the spatial utilisation. At the time of the study, the Troop 1 (green) was composed of an average of 24.9 individuals (min = 21; max = 29), Troop 2 (orange) of 19.9 individuals (18; 22), Troop 3 (pink) of 35.7 individuals (30; 40). Tracking time reached 154 days and 827 hours for Troop 1; Troop 2 = 194 tracking days, totalling 1167 hours; Troop 3 = 179 tracking days, totalling 1162 hours. The ranging areas for the three troops were 36 ha, 25 ha and 58 ha. Area estimations considered the local topography, which was extracted using the “get_elev_raster” function of the *elevatr* package (*48*) from Amazon Web Services Terrain Tiles. Houses hosting dogs are shown by the plain circles. The density of such houses was of ca. 1.66 houses/ha for Troop 1’s range, 2.14 for Troop 2, and 1.72 for Troop 3.

In 2024, Simbithi hosted 1195 domestic dogs from approximately 70 different pure breeds and additional mixed breeds (Simbithi Eco-Estate Administration, unpublished data). Dogs can be walked anywhere with or (against the estate’s guidelines) without a leash and must officially be registered. As such, their housing location is known (Figure 1). However, it does happen regularly that a dog escapes from its house and is therefore roaming freely in Simbithi, sometimes running after wildlife, vervet monkeys included. There have indeed been several reports of aggressive behaviour between dogs and monkeys, resulting in injuries to either species (Simbithi Eco-Estate Administration, unpublished data).

### Data collection

#### General data collection

The three monkey troops were tracked mostly simultaneously (52% of the tracking days included two troops, 21% included the three troops) from February 2024 to April 2025. The tracking consisted of half days (i.e., < 6 h; 61% of the troop-tracking days) or otherwise complete days (i.e., > 6 h; generally, from sunrise to sunset). The movement of the troop was assumed to be that of the human observers and was registered by use of a handheld GPS device (Blackview device, GAIA software), which was set up to record locations continuously (i.e., as soon as movement exceeding the GPS error threshold was detected).

#### Monkey–dog encounters

We recorded *ad-libitum* data of any dog-vervet monkey encounter during troop tracking. An encounter was defined when at least one dog (hereafter referred to as dogs, even if only one dog was involved) and at least one vervet monkey (hereafter referred to as monkeys, even if only one monkey was involved) were within 100 m of each other. The time and location of this encounter were recorded. We further collected information characterising the dogs’ phenotypes, the dogs’ and monkeys’ behaviours, and the context of the encounter to characterise the micro-habitat, and potential risk factors during such encounters.

We recorded the number of dogs, and the phenotype of each dog was characterised along four traits: 1) its breed, 2) its body size – small if less than 30 cm at the withers (i.e. below human knee level), or large if above, 3) its age – puppy if less than one year old (information could be later asked to the owner in case of doubt), or old if more than 10 years old, and adult otherwise, 4) its physical health condition – fit or overweight.

During an encounter, we kept track of the monkeys’ individual behaviours based on a predefined ethogram (Table S1). Because these events are unexpected and can be short, and the number of human observers is limited, we only recorded the behaviours of monkeys clearly visible within 20 m of the human observers (hence as many as possible for which we could obtain detailed and accurate data). We also kept track of the occurrence of vocalisations of all animals. Specifically, we recorded whether the dogs barked or not and collected the occurrence and type of alarm calls produced by the monkeys (chutters, barks, chirps, or unknown, following Seyfarth et al., 1980b). Finally, we characterised the monkeys’ general “troop level” reaction intensity (of all visible conspecifics) along a discrete scale: “none” when no monkeys reacted, “weak” when less than the majority did, and “strong” otherwise. We considered that monkeys reacted when they changed their behaviour due to the presence of dogs, such as being vigilant, alarm calling or moving in reaction to the dogs (see Table S1). We discarded encounters resulting from the observations of only one monkey (10%) to ensure representativeness of the troop rather than an individual.

The context of each encounter was recorded to characterise the micro-habitat surrounding the protagonists and to assess the potential risk posed to the monkeys. Specifically, we recorded whether the dogs were inside an enclosed area (e.g., a fenced garden, a car) or not, on a leash or not, and the minimal distance reached between dogs and monkeys during the encounter (less or more than 10 m, as we assumed that it represented the distance for which dogs could easily outrun monkeys), as well as the height location above ground of each monkey (less than two meters, hence a height reachable by most dogs, or higher).

### Data analyses

The objective of the study was twofold. First, we aimed to understand whether dogs were considered as a potential threat by the monkeys. Second, if so, we aimed to understand what drivers were modulating monkeys’ response intensity. To do so, we thus focused on three facets of the monkeys’ response: (1) a vocal facet, (2) an interactional facet and (3) a movement facet. All three facets should reveal antipredator responses if any, with (1) the production of alarm calls, (2) accrued vigilance, agonistic behaviour and/or immediate flee response and (3) avoidance/flee of risky areas. All data processing and statistical analyses were performed with the *R* software, version 4.3.3 (*49*).

#### (1) Vocal response

We investigated whether vervet monkeys in Simbithi elicited alarm calls as a response to encounters with dogs. We therefore fitted a binomial model regression (Model 1, Table S2), for which we considered the presence of alarm calls (i.e., whether alarm calls were emitted over the course of the encounter by at least one monkey or not) as a function of the overall intensity (weak *vs* strong, since reaction implies alarm calls) of the dog-monkey encounter. To account for potential short-term acclimation or amplification, we controlled for whether a dog encounter happened within the previous 15 min or not as a fixed effect. Because group size and demographic composition can alter anti-predator responses (Stanford, 2002), as well as influence the probability that at least one individual performs a call, we considered the troop identity as a random effect on the intercept. This term should effectively capture effects associated with troop size variation, since within-troop size variation is small and more reduced than between-troop size variation, as well as other idiosyncrasies associated with each troop.

Vervet monkeys’ alarm calls differ according to the type and intensity of the threat (*8*, *9*). Therefore, since alarm call frequency varied with the overall troop response (analysis described above, see Results), we further examined whether the vocal responses of the monkeys varied with the overall response of the troop, considered to reflect the risk perceived by the monkeys. Specifically, we measured how the frequency of each vocalisation type varied with the troop response (weak *vs* strong). We extracted 95% confidence intervals at the level of alarm call types, or alarm call types and troop, using a bootstrapping approach based on 10000 replicates (“boot” function of the *boot* package, (*50*, *51*). We assumed significant differences when the confidence intervals (“boot.ci” function) did not overlap.

#### (2) Interactional response

We used recorded behavioural data to test (1) what composed the interactional response of vervet monkeys and (2) how the perceived risk of the encounter (due to dogs’ features and the local context) influenced the monkeys’ responses.

First, we investigated whether monkeys elicited different behaviours during an encounter as a function of the intensity of the overall behavioural response, in order to identify whether an interactional tactic existed as a function of perceived risk. Here again, we fitted a binomial model regression separately for each behaviour of interest (Model 2-4, Table S2), for which we considered the general troop-level response (i.e., whether the behaviour was observed over the course of the encounter in at least one monkey or not) as a function of the overall intensity (weak *vs* strong) of the dog-monkey encounter. Specifically, we focused on three behaviours emphasising monkeys’ alertness (i.e., vigilance behaviour or looking at the dogs, see Table S1 for definitions), possible agonistic behaviour towards the dogs, or an escape behaviour (see Table S1). Again, we controlled for whether a previous encounter occurred within the previous 15 min, and the troop identity as a random factor on the intercept only. We expected that stronger reactions of the troop should be positively associated with an overall increase in the probability of alertness/escape/agonistic behaviour.

Second, we examined which labels of the dogs, and the micro-habitat context of an encounter influenced the monkeys’ general response to dogs. We focused on the general response of the troops rather than their individual behaviours for two main reasons. Firstly, the general response is more representative of the entire troop. Secondly, an interactional tactic may comprise several behaviours that are either exclusive, additive or substitutable to one another. This means that, at the behavioural level, the analysis may be noisier and/or produce more false negatives. As such, we modelled the relative probability of the troop-level response to be weak or strong (compared with no response, labelled as “none”) as a function of eight variables characterising dogs’ characteristics and the micro-habitat context of the encounter using an ordinal regression (Model 5, Table S2; “clm” function of the *ordinal* package, (*52*)). We also controlled whether at least one encounter occurred within the previous 15 minutes or not (binary variable). Only for this variable, we considered proportional odds (i.e., that the predictor estimates were the same across response levels) and otherwise considered a nominal effect for each (i.e., predictor estimates could differ across response levels) for dogs’ characteristics and the local context. Dogs’ characteristics were described by (I) their number (binary predictor: one *vs* more than one), and the risk they hypothetically represented for monkeys due to (II) their body size (binary predictor: small *vs* large), (III) their health condition (binary predictor: healthy *vs* not healthy; dogs were considered healthy if adult and fit), and (IV) if the dogs chased the monkeys (binary predictor: yes *vs* no), or (V) barked (binary predictor: yes *vs* no). The micro-habitat context of the monkey-dog encounter was described by (VI) whether the dogs could reach the monkeys (binary predictor: yes *vs* no), assuming that it was reachable if at a height lower than 2 m and at a horizontal distance of less than 10 m (these distances were taken when the monkeys reacted, or otherwise corresponded to the minimal distance between monkeys and dogs during an encounter if they did not), and (VII) the dogs’ micro-habitat (binary predictor: inside an enclosed area *vs* an open area) and (VIII) whether the dogs were on leash or not. In the cases when several dogs were present during an encounter, we considered the hierarchy of risk within the feature of interest to characterise the whole group of dogs, prioritising risky traits (being large, healthy, at a close distance, barking or chasing) over less-risky traits (being small, unhealthy, at a faraway distance, not barking or not chasing) as soon as one dog had such a risky feature. Overall, we expected the monkeys’ troop-level reaction to be more intense in case of encounters potentially perceived as risky, that is, when the dogs would be (I) numerous, (II) large, (III) healthy, (IV) chasing, (V) barking, and when the monkeys would be (VI) reachable by the dogs and (VII) when the dogs would be outside in an unfenced area.

#### (3) Movement response

Prey and predators can be engaged in a space race (*53*), inducing a space-use response of the prey to the predator presence. While, here, dogs are not hunting for monkeys, we still expected monkeys to adjust their movement according to the risk imposed by encounters with dogs, given potential lethal risk. We thus scrutinised three aspects of vervet monkeys’ movement patterns, considering the 91% of the encounters for which we had associated movement data.

First, we analysed how vervet monkeys responded to the landscape of fear posed by dogs. We distinguished between a fixed *expected* landscape of fear, shaped by the locations of dog houses, and a dynamic *experienced* landscape of fear, shaped by the locations of encounters with dogs experienced by each troop over the study period. To determine whether monkeys generally avoid areas where dogs are likely to be present, we performed an integrated step-selection analysis (*54*) with the *amt* package (*55*). We considered monkeys’ tracks at a 1-min resolution, and for each location, we sampled 50 alternative random locations. The distance and direction of these locations were respectively sampled from a Gamma distribution and a Von Mises distribution parameterised based on the step lengths and turning angles truly observed in the data. We then fitted a conditional logistic model (i.e., stratified by the “step number – Troop” id, as we fitted one model based on all three troops, Model 5), considering whether a location was chosen or not (binary response) as a function of the risk value associated with houses (continuous predictor, scaled to a mean of 0 and a standard deviation of 1 within troop), the risk value associated with encounters (continuous predictor, also scaled but within steps), and the cosine of the turning angle and the logarithm of length of a step and their interaction to account for movement advection. Risk values corresponded to the values associated with the spatial density (in probabilistic terms) associated with dog houses (expected risk) or encounters (experienced risk). The expected risk was fixed and was estimated with a Gaussian spatial kernel (“kernelUD” function of the *adehabitatHR* package, (*47*), with a smoothing parameter, *h*, of 90 m), considering all dog house locations. *A contrario*, the experienced risk was dynamic in time (*56*). To model it, we estimated daily Gaussian spatial kernels for each troop considering past encounter events (*13*). Each event was then weighted based on the time elapsed since the encounter (“sf.kde” function of the *spatialEco* package (*57*); bandwidth of the kernel set to 90 m), to mimic memory fading. We considered a maximal weight if an encounter just happened, linearly decreasing to zero at 14 days, which is longer than the reported vervet monkey short-term memory duration (*58*). We further investigated whether monkeys’ reactive behaviour was driven by spatial memory of risk. Specifically, we tested whether monkeys’ behavioural responses during dog encounters reflected a predictive representation of the spatial risk. For this, we assessed whether the magnitude of the troop’s general response (none, weak, or strong) was associated with different values of expected or experienced risk. If the monkeys had memorised these spatial landscapes of fear, we would predict them to react more vividly to coldspots than to hotspots of risk, because of potential surprise and unforeseen risk. In other words, dog encounters leading to an intense troop response should, on average, be associated with lower risk values. Conversely, no difference should be expected if the monkeys did not memorise these distributions and therefore did not anticipate the risk, or discarded risk overall. To compare risk values across the general response level of the troop (none, weak, or strong) associated with each encounter for each landscape, we used a Kruskal-Wallis test (“kruskal.test” function in the *stats* package) comparing standardised risk values to a mean of 0 and a standard deviation of 1 within each group. We then assessed the pairwise differences with Wilcoxon tests, maintaining a false discovery rate of 5% (“wilcox.test” function in the *stats* package with the *p.adjust* parameter set to “BH”).

Second, we focused on the instantaneous flight response of the monkeys. Specifically, we estimated the movement speed and the displacement (i.e., linear travelled distance) from the start of the encounter up to 15 min after it, every minute, on the raw GPS tracks rediscretised at such resolution. As the spatial data are collected by handheld GPS devices, the short-time resolution is unlikely to reflect fine-grained spatial response, but should capture long-distance flight response, if any. Using log-Gaussian generalised linear mixed models (Model 6 and 7, Table S2), we investigated how these two metrics varied as a function of the general troop response and the time elapsed since the encounter (scaled to a mean of 0 and a standard deviation of 1, to minimise convergence problems), considered as a polynomial term of second order to allow for some plateauing or transient responses, and set in interaction with the general troop response. We also considered, as before, whether a previous dog encounter occurred within the previous 15 min. We accounted for temporal autocorrelation using an autoregressive process of first order (“ar1” function) and included the troop identity as a random effect on the intercept. This formed the *full* model, while the *null* model (to which we compared the *full* model) discarded the general response of the troop. We fitted the models with the “glmmTMB” function of the *glmmTMB* package (*59*). A sharper increase in the speed and the displacement over time, with a more intense general response compared to the absence of response, would evidence a fleeing strategy, while the opposite would suggest a freeze strategy.

Finally, we focused on the subsequent avoidance behaviour of the monkeys. Specifically, we computed the first passage time since the encounters using the *recurse* package (*60*). We considered a circular buffer of 50 m around the encounter location and estimated the time delay for monkeys to revisit this circular area after the encounter. Note that we considered a revisit only if the monkeys had left the area for more than 20 min. We then investigated how the first passage time varied as a function of the general troop response using Cox Proportional-Hazards Model (Model 9, Table S2; fitted with the “coxph” function of the *survival* package, (*61*, *62*)). In this model, we also considered the logarithm of the number of days the monkeys were not tracked up to the first passage time as a fixed predictor, to control for possible missed recursions, and the troop identity as a random effect on the intercept (“frailty” function of the *survival* package, (*61*, *62*)).

#### (4) Statistical implementation

For all generalised linear mixed models (inventory in Table S2), prior to testing for any singular effect for each analysis (comparing the likelihood of models with and without the variable of interest), we ensured that the tested variables (hence, if several) significantly affected the response together by comparing the likelihood of the *full* model (containing all fixed and random predictors) to that of a *null* model (containing only control predictors, including random ones), using a likelihood ratio test (“anova” function of the *stats* package readily available in *R*). Furthermore, we ensured that there was no problem of collinearity using the Variance Inflation Factor (VIF, all VIFs were inferior to 2; “check_collinearity” function of the *performance* package, (*63*)). We also proceeded to further model diagnostics adapted to each model we fitted (see Supplementary Material, Model Verifications; Figures S1-8).

We discuss the models in the results section based on the marginal effects (i.e., when the effects of all other covariates are averaged) and their associated 95% confidence intervals extracted with the *emmeans* package, (*64*) or the *ggeffects* package, (*65*), depending on the model type. Complete model outputs are available in the Supplementary Material, Tables S3-6. For all regressions, we then estimated pairwise comparisons (relative probabilities, odds ratios - OR, etc.) and their associated confidence intervals based on their marginal effects, using a Monte-Carlo approach, assuming a Gaussian distribution of the estimates.

### Ethical statement

Data sampling was non-invasive. The research was approved by the board of Simbithi Eco-Estate and the ethics committee of the University of KwaZulu Natal (AREC/00004821).

## Results

Over 14 months, the three studied troops of vervet monkeys encountered domestic dogs 843 times (Troop 1: 191 times, Troop 2: 284 times and Troop 3: 368 times). 755 of these encounters (90%) included more than one monkey. Among those, 516 (68%) included the focal individuals reacting to the dogs. Yet, within a troop, the response was not necessarily uniform, and in 30% of the time for which at least one monkey reacted, at least another one did not.

### Vocal response

Alarm calls happened in 55% of the 755 analysed encounters. In these cases, it most frequently (86%) included chutters, while barks and chirps were rare (< 8%, Figure 2).

**Figure 2.**
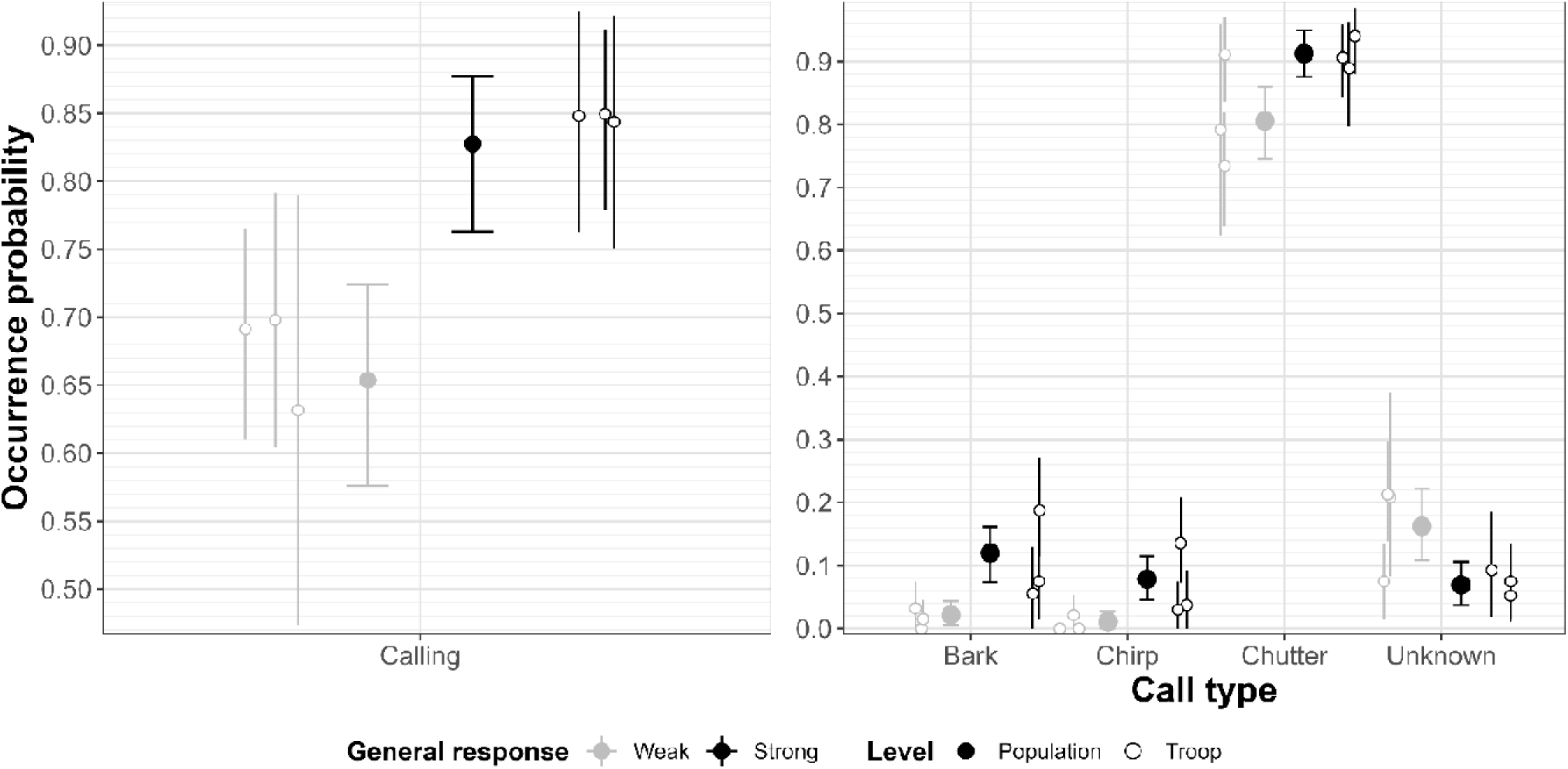
Vocal response of vervet monkeys to dog encounters. **(Left) Probability of alarm calls to occur at the troop level as a function of the troop’s general response level** Alarm calls are more likely to occur when the troop reacts strongly. The plain circles represent the model marginal estimates, while the segments represent the associated 95% confidence intervals. (Right) Occurrence rate of the three types of alarm calls (and when unidentified) as a function of the troop’s general response level (weak *vs* strong). The occurrence of alarm calls varied with the level of general response of the troop. The circles represent the overall population (plain) or troop (open) averages, and the vertical segments represent the associated 95% confidence intervals obtained using a bootstrapping approach. Note that those are conditional probabilities when at least one call occurred. Furthermore, several calls can co-occur.

The odds of alarm calls happening were 2.66 times [1.52, 4.54]_CI95%_ higher when the troop strongly reacted compared to when it only weakly did (Figure 2). Furthermore, alarm call types differed with the general response of the troops. In particular, barks and chirps were respectively 7.34 [2.34, 23.87]_CI95%_ and 8.34 [2.34, 19.18]_CI95%_ times more likely to occur at the troop-level when the troop strongly reacted compared to when it weakly did (Figure 2). Chutters occurrence was almost unaffected by whether the troop strongly or weakly reacted (increase of 1.13 [1.04, 1.23]_CI95%_, Figure 2).

### Interactional response

Monkeys displayed different behaviours according to the troop’s general response. Monkeys were mostly alert (> 80% of the time, Figure 3) in both weak and strong troop-level reaction (χ²_1_ = 2.075, p = 0.150). The odds of agonistic behaviours occurring (from the monkeys towards the dogs) were 4.44 [1.09, 8.82]_CI95%_ times higher when the troop reacted strongly compared to when it only reacted weakly (Figure 3). The odds of fleeing were also 3.90 [1.92, 7.69]_CI95%_ times higher when the troop reacted strongly compared to when it only reacted weakly (Figure 3).

**Figure 3.**
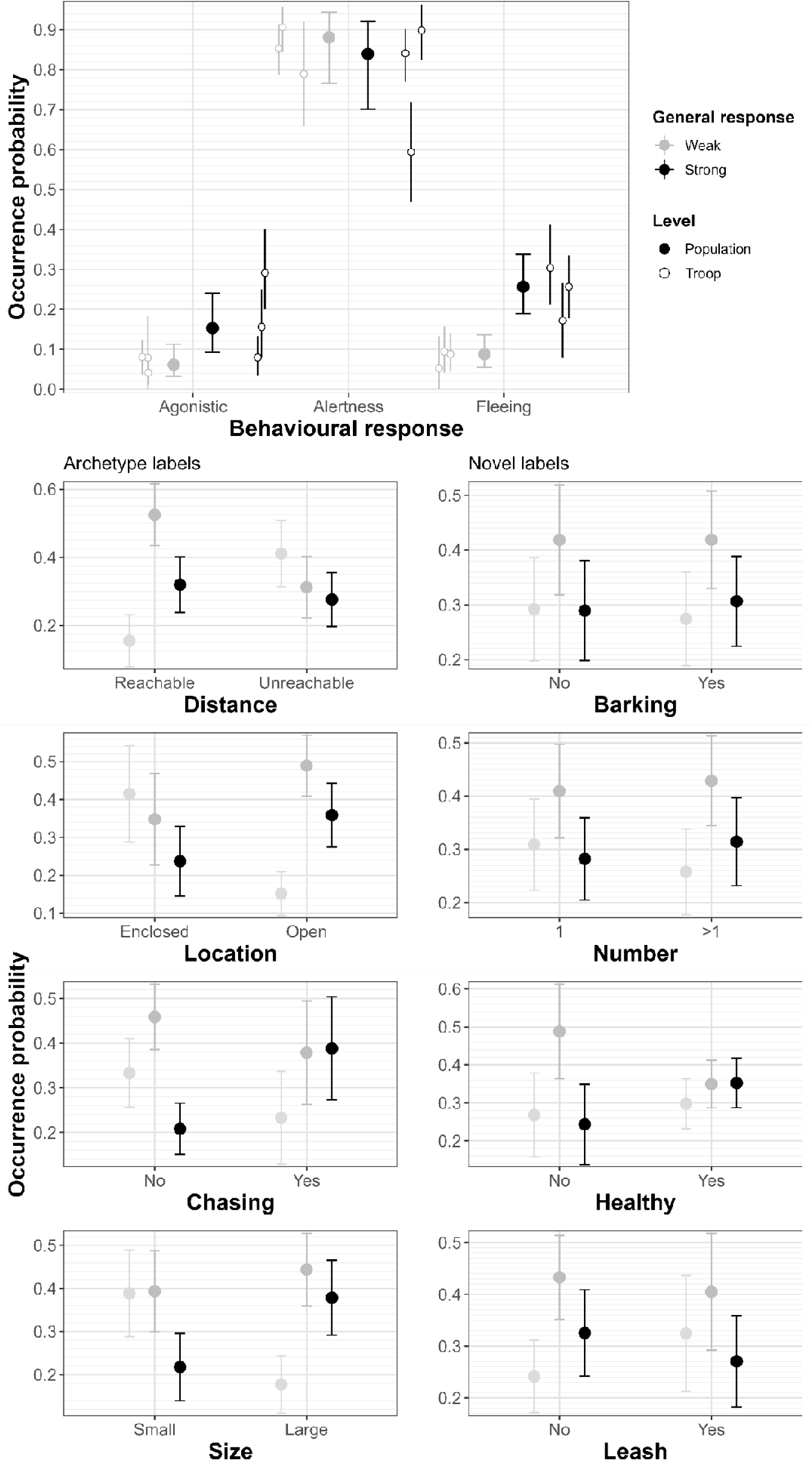
Interactional response of vervet monkeys to dog encounters. **(Top panel) Behaviour occurrences as a function of the troop’s general response.** Alertness state is highly likely in the case of dog encounters. Strong behavioural response of the troop is associated with increased agonistic and fleeing behaviours. The open circles represent the troop-level observed estimation, while the segments represent the associated 95% confidence intervals. They were estimated using a Monte-Carlo approach, assuming a Gaussian distribution of the estimates. The plain circles represent the model marginal estimates, while the segments represent the associated 95% confidence intervals. **(Bottom panels) Troop general response level as a function of dogs’ features and encounter micro-habitat context.** Vervet monkeys respond more strongly to traditional, but not novel, features of risk. The plain circles represent the model marginal estimates, while the segments represent the associated 95% confidence intervals.

Overall, the context of the encounter and the dogs’ features influenced the monkey troops’ level of reaction (*full vs null* model likelihood ratio test: χ²_17_ = 138.070, p < 0.001). This was primarily because of the dogs’ size, whether the monkeys were reachable by the dogs or not, and due to the dogs’ behaviours, specifically whether they chased them or not (Figure 3). The odds were 2.32 [1.23, 4.25]_CI95%_ times to react strongly when dogs were large than when they were small. When dogs could reach them because they were at a close distance (both horizontally and vertically) than when they were not, the odds were 2.55 [1.40, 4.45]_CI95%_ times higher for monkeys to weakly react, and 1.28 [0.71, 2.20]_CI95%_ times to strongly react, respectively. Similarly, when the dogs were located in an open area, compared to when dogs were in an enclosed area, the odds were 1.92 [0.97, 3.58]_CI95%_ times and 1.93 [0.97, 3.67]_CI95%_ times higher to weakly or strongly react, respectively. Finally, when dogs chased after them, compared to when they did not, the odds were 2.54 [1.30, 4.50]_CI95%_ times to strongly react. Conversely, there was weak evidence that the number of dogs (χ²_2_ = 2.976, p = 0.317), the dogs’ health condition (χ²_2_ = 5.626, p = 0.060), barks emitted (χ²_2_ = 0.176, p = 0.916), or whether the dogs were on leash or not (χ²_2_ = 2.748, p = 0.253) influenced the level of reaction of the troop.

### Movement response

The three monkey troops tended to use areas surrounding houses where dogs were present – representing the expected landscape of fear – more than would be randomly expected, with the odds of choosing a location increasing by 8% with every standard deviation increase in associated risk (est [CI_95%_] = 1.08 [1.03, 1.13]; Table S5). Conversely, they did not consider previous encounters (i.e., the experienced landscape of fear) in their movement choices (est [CI_95%_] = 1.00 [0.98, 1.02]). Additionally, the troop’s general response intensity depended on the expected landscape of fear but not the experienced landscape of fear (Kruskal-Wallis tests: p = 0.008 and p = 0.064, respectively, Figure 4A). Specifically, the expected risk was 1.87 [1.09, 4]_CI95%_ times lower when they strongly reacted compared to when they did not react (Figure 4A).

**Figure 4.**
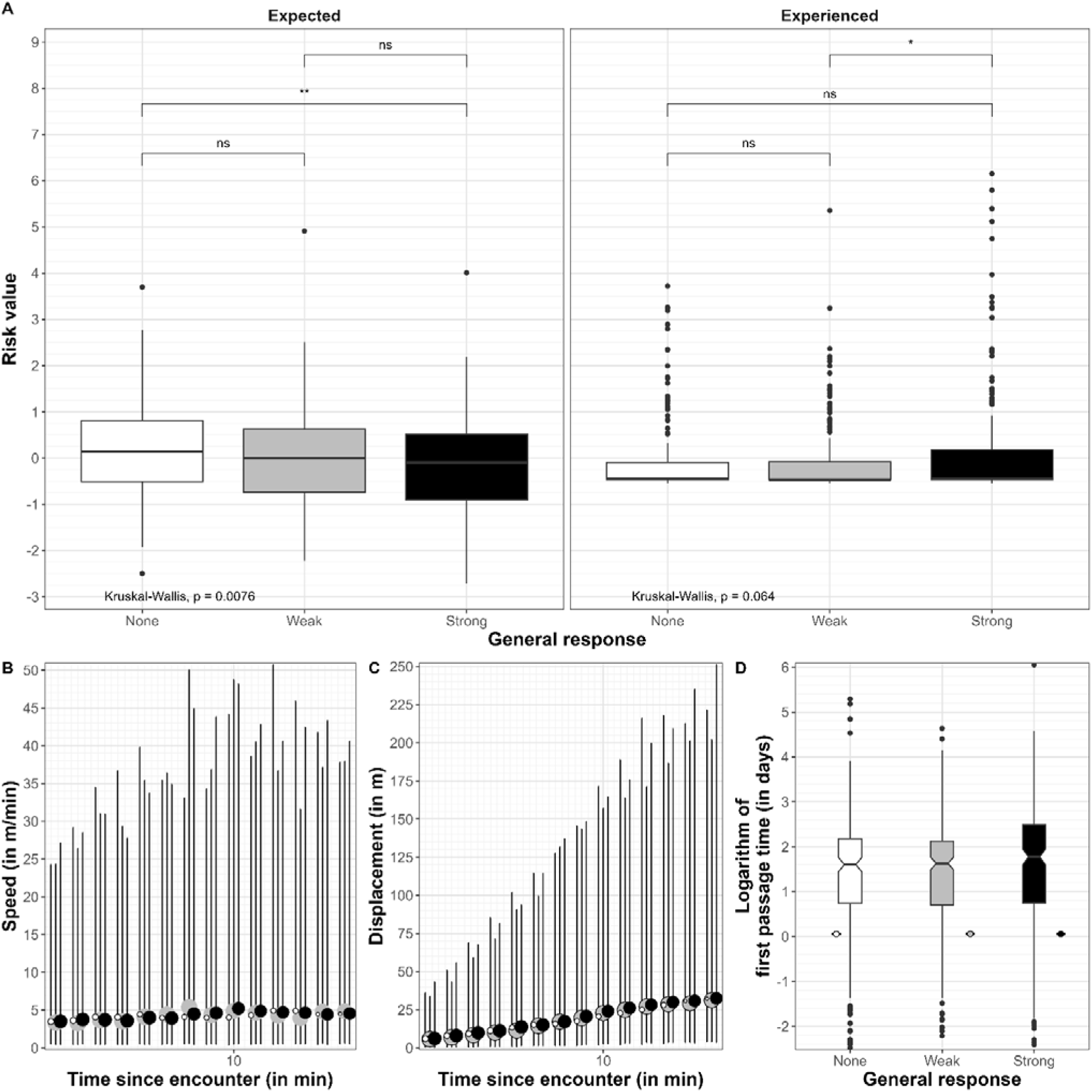
Movement responses of vervet monkeys to dog encounters. **(A) Vervet monkeys reacted more vividly when risk was unexpected.** The observed risk values at each location where dogs were encountered are represented by classical boxplots, according to the standardised (within troop) expected or experienced landscape of fear. Colours indicate the troop’s general response level (white: no response; grey: weak response; black: strong response). The statistical significance of the differences between the three levels or between pairs of levels is indicated at the very bottom or top, respectively, and was assessed using a Kruskal–Wallis test or Wilcoxon tests, respectively. *: p < 0.05, **: p < 0.01, ***: p < 0.001, ns = not significant. **(B, C) Travel distance and displacement (i.e., beeline distance) as a function of the general response level.** Monkeys travelled and displaced similarly in the 15 minutes following an encounter with dogs, independent of how the troop reacted. The observed data are shown by the plain circles, which were dodged for readability. The model predictions and the associated 95% confidence intervals are shown by the plain line and the shaded polygons. **(D) First passage time to past dog encounter locations as a function of the troop general response level.** The time monkeys took to return to the locations of previous encounters with dogs is insensitive to whether they weakly or strongly reacted. The observed data are represented by the notched boxplots. The model predictions are added on the side of each, with plain circles representing the mean, and the segments their associated 95% confidence intervals.

Whether the troop did not react, reacted weakly or reacted strongly to an encounter had no consequences on the overall troop’s subsequent movement speed and displacement (*full vs null* model likelihood comparisons: χ²_6_ = 4.444, p = 0.616 and χ²_6_ = 3.691, p = 0.718, respectively, Figure 4B and C).

Finally, after dog encounters, the return time to the encounter location was independent of the reaction level the troop exhibited during the encounter (*full vs null* likelihood ratio test: χ²_2_ = 2.136, p = 0.349, Figure 4D).

## Discussion

In ever-changing environments, animals must constantly adapt and adjust their tactics to maintain high fitness. In a prey-predator context specifically, successful behavioural adaptation by the prey is necessary for survival but sometimes changes related to threat exposure happen faster than traditional evolutionary timescales (*66*). This is particularly true in the case of human-caused habitat transformation, questioning the ability of wild animals to recognise and cope with new predators (*23*). In this study, we investigated how vervet monkeys coped with a novel predation risk posed by the cohabitation with domestic dogs in an anthropised habitat. We demonstrated that the vervet monkeys from this semi-urban ecosystem elicited an ingrained predatory response to the archetype labels of risk of this novel threat but failed to discriminate the new cues of this evolutionary novel danger.

### Vervet monkeys show a fear response to dogs

In response to the presence of dogs, vervet monkeys showed strong fear reactions, suggesting that they consider dogs to be a threat and are thus not completely naïve (*28*). Dogs are indeed phylogenetically close to some of the vervet monkeys’ natural predators. In addition, while they have not been reported as predators to our knowledge, vervet monkeys live in some areas in sympatry with wild canid species, such as wild dogs or jackals. This may facilitate the recognition of certain risk labels (*23*, *67*). Consequently, the urbanised vervet monkeys responded to this threat mostly similarly to real predators in their natural habitat.

Vocally, they emitted alarm calls, and quite trivially did so more frequently when the whole troop reacted more strongly. Alarm calls enable the transmission of information about the threat to all members of the troop, triggering defensive behaviour (e.g., climbing up in case of terrestrial danger). Although these calls are usually associated with a specific type of predator, we observed some plasticity: these monkeys used both alarm-call types usually produced towards terrestrial and aerial predators, when encountering dogs (*8*, *9*). This strengthens the point that these calls may more encode the threat level (i.e. a quantitative trait) and not only the predator type (i.e. a qualitative trait) (*68*, *69*). In this case, it appears quite necessary, as for such a “novel” predator, specific calls do not yet exist. Yet, they may be developed in the future by combining existing calls. Indeed, it was recently shown that primates can combine existing calls to create new ones (*70*).

Interactionally, as expected, vervet monkeys increased their agonistic responses and attempts to flee when the troop reacted more, that is, when they considered being threatened. Interestingly, agonistic behaviours (from the monkeys towards the dogs) were observed as frequently as fleeing behaviours. Although rare, cases have been reported at Simbithi in which the monkeys injured the dogs too. When prey-predator asymmetry is not considerable, prey may indeed retaliate rather than flee (*14*). This suggests that vervet monkeys may consider dogs to be potentially risky, but not necessarily deadly. This happens with Phayre’s leaf monkeys in India too, which also retaliate against domestic dogs (*42*). The threat the monkeys perceived appeared to be calibrated by specific labels associated not only with the dog, but the context of the encounter (see **Vervet monkeys fail to discriminate novel labels of risk**).

From a movement perspective, vervet monkeys showed proactive, rather than reactive, responses. Yet, as opposed to traditionally observed in the wild (*71*), this proactive behaviour consisted of a risk-prone behaviour, with monkeys exploiting preferentially areas associated with dog houses. Yet, the monkeys’ reaction to dog encounters was more vivid in the coldspots of the expected risk (i.e., the monkeys reacted strongly when the risk due to dog-house presence was weak, hence in areas of unexpected risk). This follows the violation-of-expectation paradigm (*72*), in which a stronger behavioural response is expected when a situation deviates from what was anticipated, and thus suggests that this risk-prone behaviour is deliberate. In other words, this illustrates that monkeys were somehow surprised when a dog was encountered while they expected low risk. In Simbithi, houses are associated with significant food resources, which likely overweigh the risk of encountering dogs, hence a certain attractivity despite risk for those houses hosting dogs. Houses are indeed generally surrounded by gardens with fruit trees and shrubs that monkeys feed on. Furthermore, open windows and doors may provide monkeys with access to human food inside the house (*45*). As vervet monkeys are known to possess efficient spatial cognition (*73*, *74*), they likely mentally “mapped” the dog houses, and as such can recognise, and anticipate, risky areas. Yet, dog houses are not necessarily the sole locations where dogs can be met. In this regard, we found no evidence of monkeys keeping track of a dynamic landscape of fear, based on past experience of dog encounters: the spatial distribution of risk associated with previous encounters (i.e., the experienced landscape of fear) neither impacted movement choices nor the intensity with which monkeys reacted. Overall, these results tend to emphasise that monkeys do not respond to risk *per se*, but rather to how the perceived risk aligns with long-term expectations.

When risk was faced nonetheless, aside from immediate flight over a few meters, particularly climbing up, we observed no reactive long-distance movement responses within the 15 minutes following an encounter (Figure 4). The monkeys did not travel faster or move away more if the perceived threat changed. This absence of reactive movement was also observed in the wild, where encounters with natural predators did not affect subsequent movement patterns (*75*). Living under permanent fear, it is possible that threat encounters only induce an increase in subsequent short-term alertness, which we did not monitor thoroughly after the encounter happened. Such short-term interactional, but not movement, reactive response appears particularly relevant in this habitat, since dogs are highly prevalent and the number of encounters largely surpasses the usual amount with natural predators. As such, encounters are inevitable. Maintaining a high travelling speed, or important advective movement, would certainly be too energetically costly in this case, and may further induce a lot of lost foraging opportunities. Furthermore, being arboreal too, vervet monkeys may simply seek refuges by climbing up trees, and not by moving horizontally away. Such a fine-grained response could not be captured by our data collection, as it was based on handheld GPS devices carried out by human observers, hence likely reflecting more the “human behaviour” rather than the monkeys’ one at fine resolution. However, this fits coherently with the observed increase in immediate flight behaviour when the troop strongly reacted (Figure 3, top). In fact, 73% of the encounters happened in a local environment where nearby vertical escape was possible (e.g., presence of close trees, fences, etc.). In addition, monkeys did not take longer to return to locations where threatening encounters occurred than when encounters were calm. Possibly, recursions to a location may be more strongly driven by resource availability than by predation risk. Vervet monkeys are capable of tracking spatiotemporal cues (*76*), which are associated with the pulsating regime of their resources, the main driver of most animals’ recursions (*77*, *78*). However, their behaviour during visits (e.g., alertness) may possibly vary as a consequence of previously perceived threats, as typically occurs in prey (*79*), and this remains to be explored.

### Vervet monkeys fail to discriminate novel labels of risk

Since responding to a threat incurs costs, we expected vervet monkeys to modulate their fear response according to the risks of dog encounters. To this point, we demonstrated that vervet monkeys consider labels of hazard and exposure similar to those they have faced throughout their evolutionary history (e.g., escapability at the current location as a function of exposure/vulnerability within the habitat and distance to the predator, predator size, and chasing behaviour). This shows again that vervet monkeys are not completely naïve and use archetypal labels that highlight some risk of their natural predators to modulate their response to predation risk (*80*). Thus, their response to dogs likely reflects ingrained antipredator behaviour that usually does not disappear in prey even if their natural predators are absent from the area (*81*, *82*).

Nonetheless, vervet monkeys failed to discriminate levels of risk based on novel labels, such as whether the dogs barked, were in a healthy condition, or were on a leash. This finding suggests that the monkeys may find these labels irrelevant, may not yet have learned the meaning of these labels, or alternatively, do not have the cognitive tool kit in place to comprehend such distinctions. Surprisingly, we found that this was the case for the number of dogs, which was not a primary driver of the monkey response, yet some primates can assess quantity (*83*). The absence of quantitative judgment in our data may indicate that they do indeed apply hierarchical reasoning, prioritising the riskiest scenario (i.e., the riskiest dog labels), hence discarding dog number. However, this could also be a statistical artefact. Indeed, when a hierarchical rule is applied to associate one label with a group of dogs, the larger the number of dogs, the more likely it is that one dog will have a risky label. This means that the effect of the number of dogs may be diluted by the inherent (even if weak) collinearity between dog number and the other risk labels, leading to a statistical artefact of absence of significance. To verify this, we built a model based on a majority rule, which does not create such a collinearity. Again, it did not identify dog number as a relevant variable either (see Figure S8, Table S7; see Supplementary Material **Statistical artefact for the variable “dog number”?**), suggesting that our conclusions are biologically meaningful rather than a statistical artefact.

Finally, outside of urban settings, labels incorporating human artefacts such as leash condition are actually irrelevant, which likely explains the absence of response to it, at least for now. Although the urban environment may not transform a species’ cognitive capacities per se (*84*), it is plausible that these response patterns will change over evolutionary time, should the species’ cognitive tool kit be flexible enough, and the selective gradient be strong enough to select for learning. The latter requires additional data on dog-related mortality to evaluate the proportion of mortality caused by dogs. If this is substantial, an adaptive antipredator response would still generally require considerable learning and causal understanding, and even more so with species that show distant phylogenetic relationships with native predators (*67*, *85*). As the eco-estate is only a few decades old, this may have not happened yet, but we can suppose that, owing to their advanced cognitive skills (*86*), vervet monkeys may progressively acquire this knowledge. Magpies (*Grallina cyanoleuca*), which are cognitively well-endowed and have lived in close proximity to dogs for a long time, did eventually react differently to the presence of dogs depending on whether they were on a leash or not (*87*): they maintained a flight response but only displaced walking when the dog is on a leash, and flew when it is not, suggesting that they understood that a leash can restrain the dog’s movements (*87*). In addition, it is important to note that a lack of reaction does not necessarily indicate a lack of comprehension. Barks are generally a sign of aggression, fear or excitement in dogs (*88*). As such, howler monkeys (*Alouatta palliata*) innately reacted with increased vigilance and stress hormones in response to bark playback experiments (*89*). However, contrary to playbacks, real barks also reveal the presence of dogs. Once spotted, the level of risk they represent as terrestrial predators in an urban environment (where climbing facilities are plentiful) may be low in reality. This could explain why, in our context, monkeys did not exhibit increased responsiveness to barking dogs.

## Conclusions

In this study, we examined the interactions between vervet monkeys and domestic dogs in Simbithi Eco-Estate, Ballito, South Africa. In response to encounters with dogs, vervet monkeys exhibited antipredator behaviours, including alarm calling, increased vigilance, and agonistic reactions, but deliberately engaged in risky areas based on spatial memory and risk anticipation. Novel cues of risk, whether related to dog morphology (e.g., physical health condition) or the human-altered context (e.g., presence or absence of a leash), did not influence the intensity of their responses. This contrasts with their reactions to honest labels of risk also associated with predation in a “natural” context (e.g., distance, etc.), which did modulate response intensity. These findings suggest that vervet monkeys’ responses to evolutionarily new predators such as domestic dogs are ingrained. Whether this apparent rigidity in vervet monkeys’ antipredator tactic is due to an inability to learn new cues, to low fitness incentives (e.g., if mortality in this population is more often caused by vehicle collisions than by dogs’ attacks), or ongoing learning and adaptation yet not visible remains an open question.

## Acknowledgements

We thank Margi and Derrick Lilienfeld, Ayanda Duma and the entire Simbithi board for their support of our research. We further thank all students and researchers who joined the Urban Vervet Project for contributing to data collection during the time of the study. We thank Mathilde Martin for discussing several points of the study. This work was funded by the KONE Foundation (Finland), the Swiss National Science Foundation (Switzerland), Grant Nrs CRSK-3_220769 and PZ00P3_202052, and the University of Zurich’s FAN Förderung der Akademische Nachwuchses (die Gebauer Stiftung, Switzerland). CV was funded by the European Union - NextGenerationEU, under the National Recovery and Resilience Plan (NRRP), Project title “National Biodiversity Future Center -NBFC” (project code CN_00000033, CUP J33C22001190001). Views and opinions expressed are, however, those of the authors only and do not necessarily reflect those of the European Union or the European Commission, nor can they be held responsible for them.

## Data availability

The data will be archived on a public repository upon publication, together with the codes used to compute analyses. The codes are also available at https://github.com/benjaminrobira/VervetAndDogs.

## Author contributions

BR: conceptualization, data curation, formal analysis, software, supervision, methodology, visualization, writing – original draft preparation, writing – review & editing

NB: conceptualization, data curation, investigation, methodology, writing – review & editing

SM: conceptualization, data curation, investigation, methodology, project administration, supervision, writing – review & editing

AM: validation, writing – review & editing CV: writing – review & editing

SF: conceptualization, funding acquisition, project administration, writing – review & editing

## Supplementary material

### Behavioural Ethogram

**Table S1.** Ethogram used to encode monkeys’ behaviours based on *ad-libitum* sampling during each monkey-dog encounter.

| Behaviour name | Behaviour Category | Behaviour Definition |
| --- | --- | --- |
| Biting | Agonistic | The individual puts its open mouth in physical contact with any body part of the dog |
| Chasing | Agonistic | The individual approaches the dog at high speed to chase it away. |
| Head bobbing | Agonistic | The individual stares at the dog and does some approach/retreat movement with his/her head. |
| Staring | Agonistic | The individual is looking at the dog in an aggressive way, repetitively raising eyelids. |
| Focusing/Looking | Alertness | The individual is watching the dog by stopping its current behaviour, either continuously for at least 3 sec (focusing) or from time to time (looking). |
| Vigilance | Alertness | The individual stares at the dog for quite a long time, and is being vigilant with a straight back, high head, and can even be bipedal. |
| Fleeing | Escaping | The individual runs away/climbs up at high speed from the dog. |
| Alarm calling | Vocal | The individual produces alarm calls towards the dog. Alarm call types are chutters, barks or chirps. |

### Model Outputs

We present below a series of tables obtained with the “tab_model” function of the *sjPlot* package (*90*) (generalised linear models, proportional hazard model) or manually directly from the model output (“tidy” function from the *broom* package, (*91*), for the ordinal model), and presenting the raw model outputs (see Table S2 for an inventory of the models).

**Table S2.**
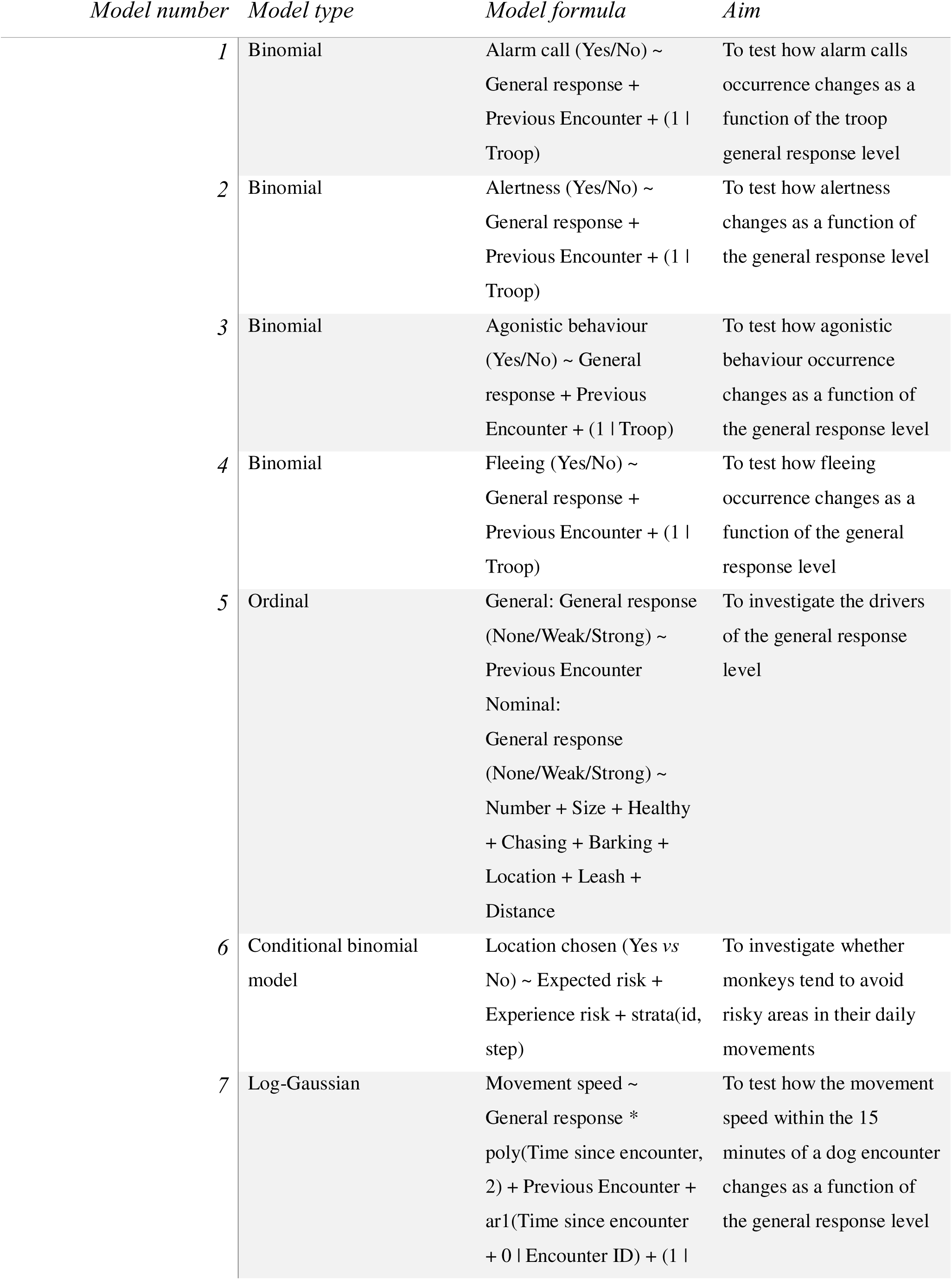
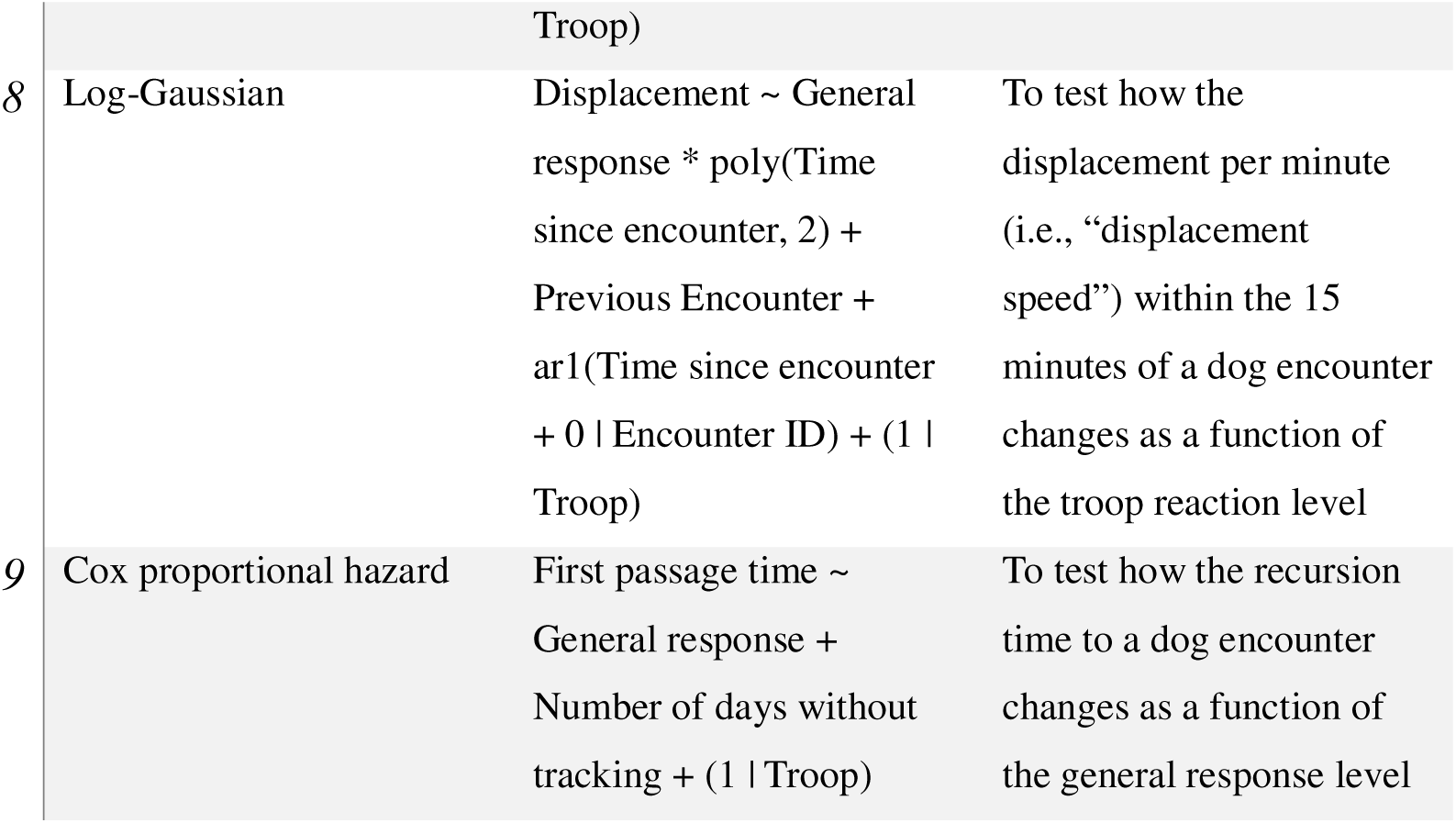
Inventory of the statistical models used in this study. The model formula is written in the *R* programming language.

**Table S3.**
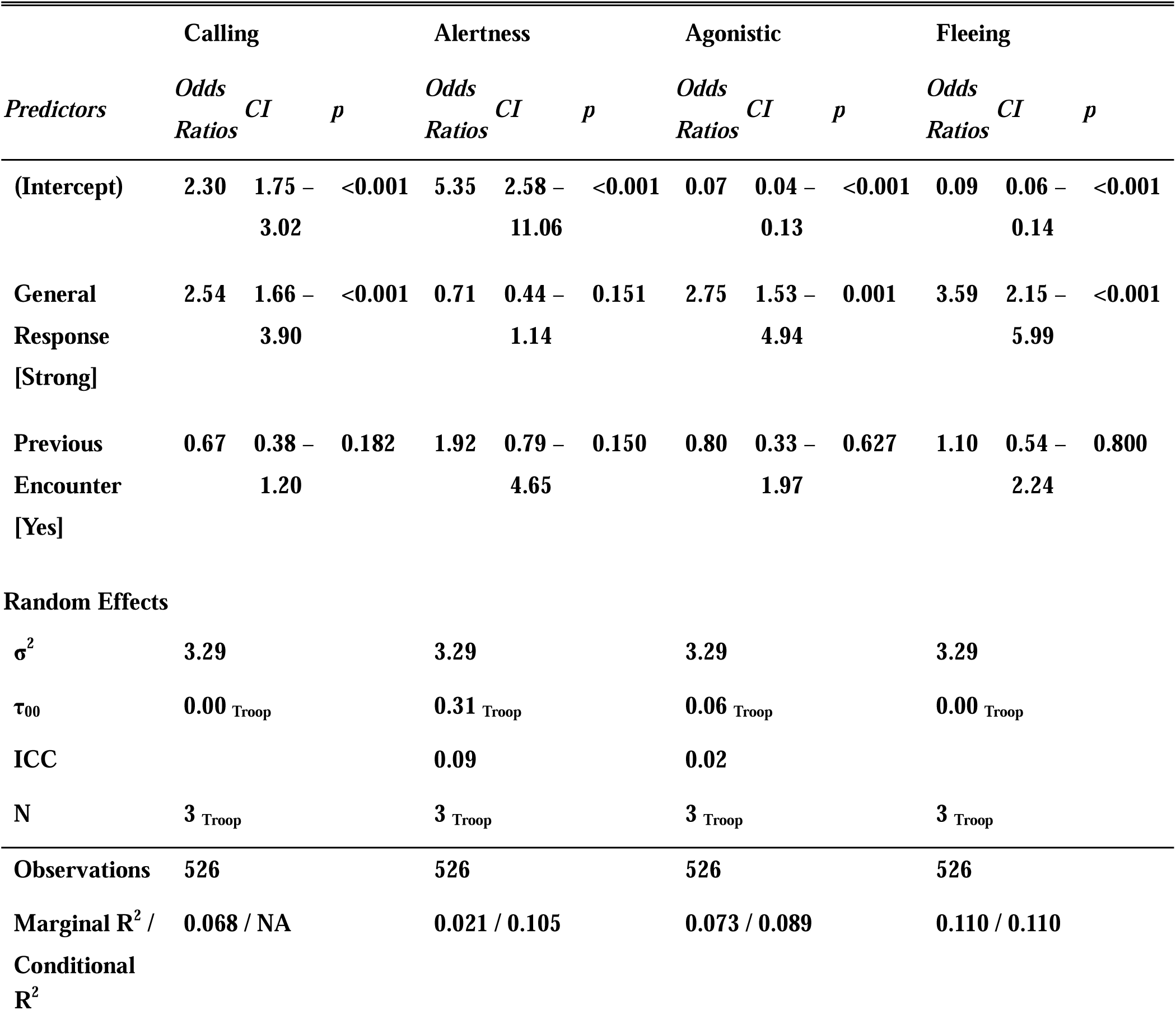
Output of the model 1, 2, 3, 4 testing the variation of types of behaviours as a function of the troop response level. CI = 95% confidence interval, *p* = p-value.

**Table S4.** Output of the model 5 investigating the drivers of the troop general response level.

| Level | Term | Estimate | Std.error | Statistic | p-value |
| --- | --- | --- | --- | --- | --- |
| None\|Weak | (Intercept) | -0.536 | 0.505 | -1.062 | 0.288 |
| **Weak\|Strong** | **(Intercept)** | **2.239** | **0.450** | **4.977** | **0.000** |
| None\|Weak | Number [>1] | -0.348 | 0.236 | -1.473 | 0.141 |
| Weak\|Strong | Number [>1] | -0.173 | 0.199 | -0.870 | 0.384 |
| **None\|Weak** | **Size [Large]** | **-1.398** | **0.244** | **-5.738** | **0.000** |
| **Weak\|Strong** | **Size [Large]** | **-0.858** | **0.236** | **-3.630** | **0.000** |
| None\|Weak | Healthy [Yes] | 0.201 | 0.355 | 0.565 | 0.572 |
| Weak\|Strong | Healthy [Yes] | -0.585 | 0.305 | -1.917 | 0.055 |
| None\|Weak | Chasing [Yes] | -0.674 | 0.364 | -1.851 | 0.064 |
| **Weak\|Strong** | **Chasing [Yes]** | **-0.958** | **0.281** | **-3.404** | **0.001** |
| None\|Weak | Barking [Yes] | -0.118 | 0.337 | -0.351 | 0.725 |
| Weak\|Strong | Barking [Yes] | -0.093 | 0.274 | -0.340 | 0.734 |
| **None\|Weak** | **Location [Open]** | **-1.721** | **0.398** | **-4.321** | **0.000** |
| **Weak\|Strong** | **Location [Open]** | **-0.653** | **0.302** | **-2.160** | **0.031** |
| None\|Weak | Leash [Yes] | 0.560 | 0.377 | 1.485 | 0.138 |
| Weak\|Strong | Leash [Yes] | 0.296 | 0.268 | 1.102 | 0.270 |
| **None\|Weak** | **Distance [Unreachable]** | **1.677** | **0.302** | **5.562** | **0.000** |
| Weak\|Strong | Distance [Unreachable] | 0.235 | 0.204 | 1.152 | 0.249 |
|  | PreviousEncounter [Yes] | -0.273 | 0.248 | -1.101 | 0.271 |

**Table S5.**
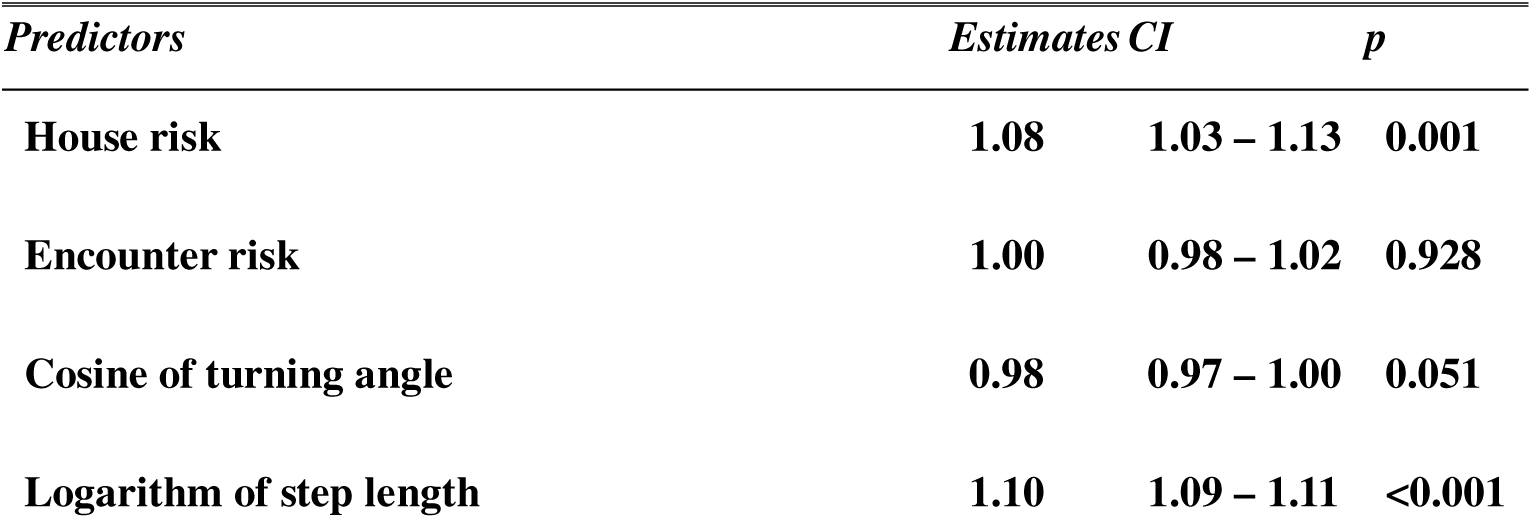
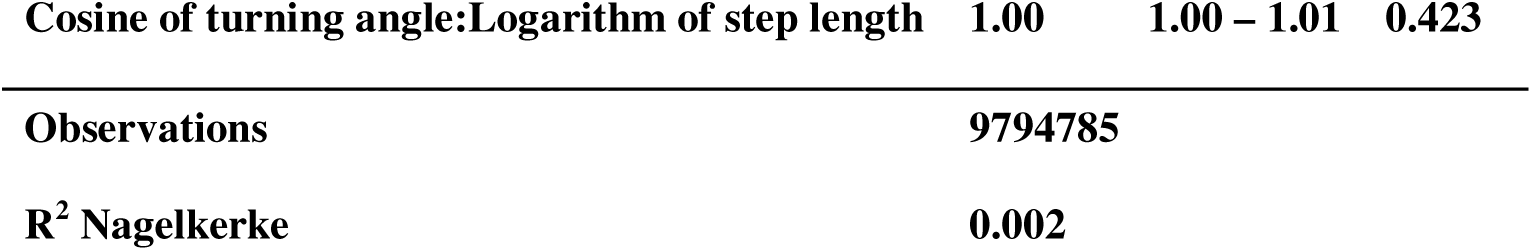
Output of the model 5 testing whether vervet monkeys consider expected/experienced risk in their daily movement choices (step-selection analysis framework). CI = 95% confidence interval, *p* = p-value. The estimates were exponentiated.

**Table S6.**
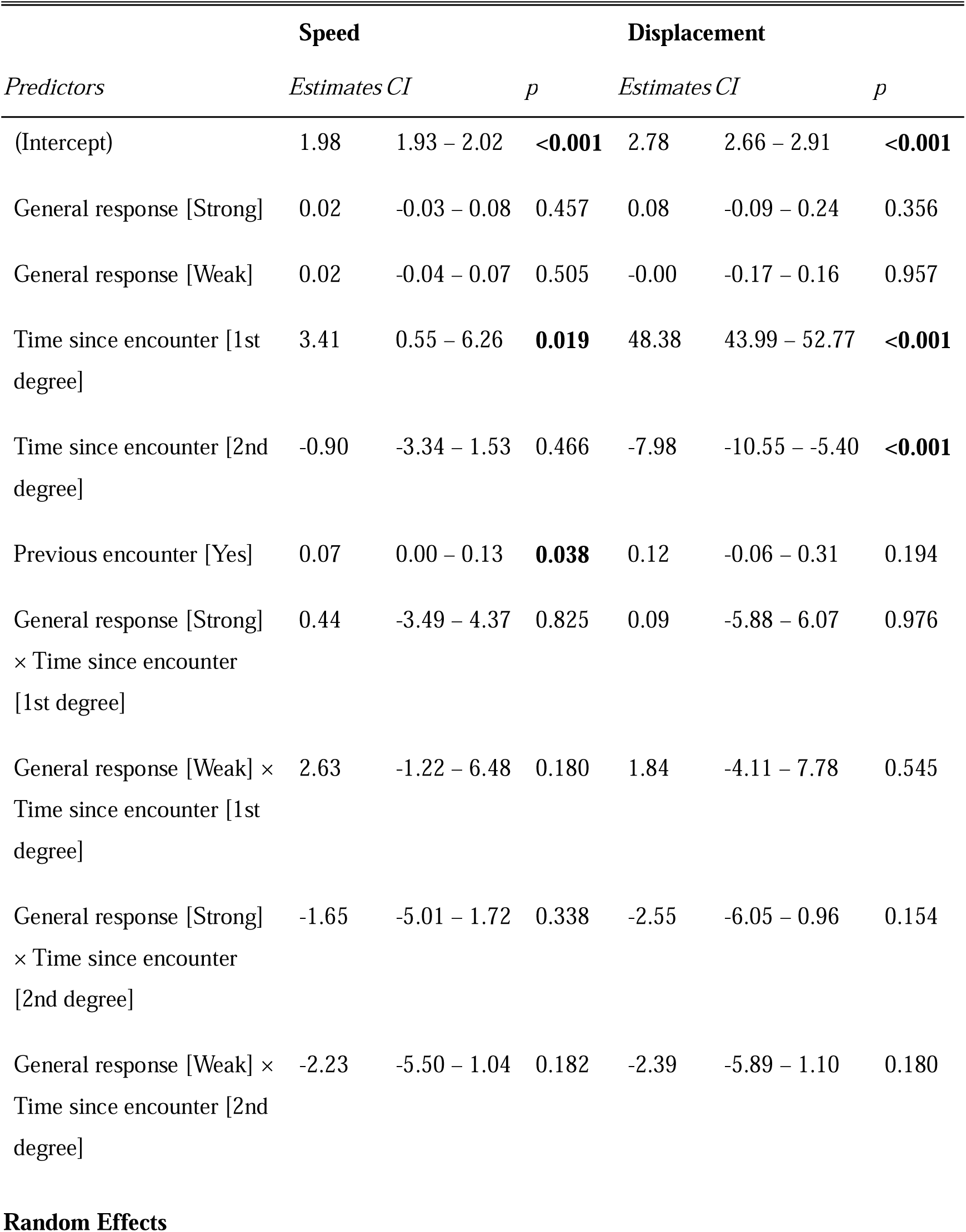
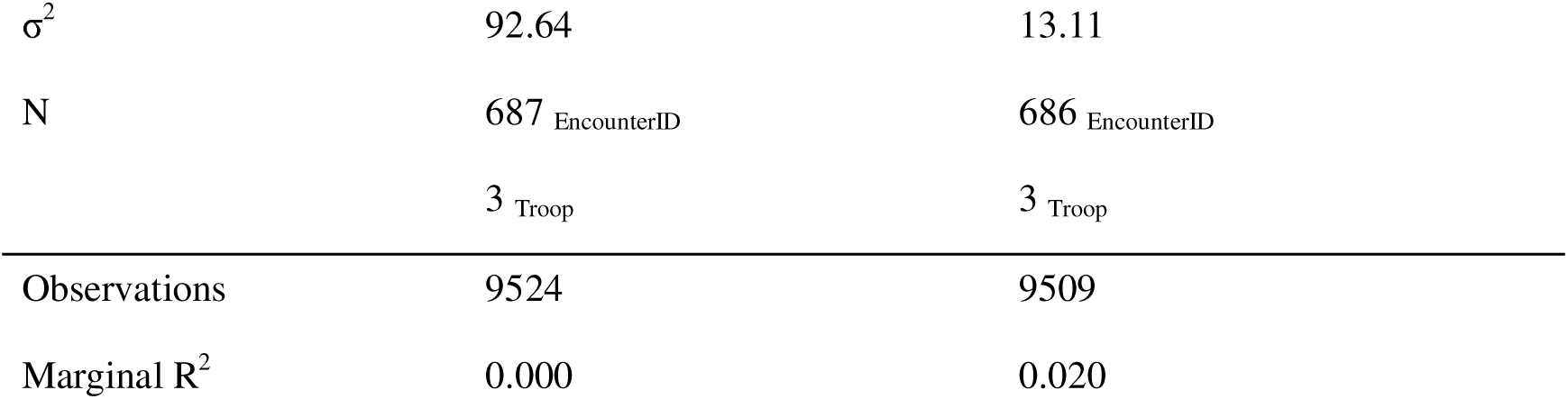
Output of the model 7 and 8 testing the variation in speed (in m/min) and displacement (in m) as a function of the troop’s general reaction level and time elapsed since the encounter (in min; max 60 min). CI = 95% confidence interval, *p* = p-value. The estimates were exponentiated.

| *Predictors* | **Speed** |  |  | **Displacement** |  |  |
| --- | --- | --- | --- | --- | --- | --- |
|  | *Estimates* | *CI* | *p* | *Estimates* | *CI* | *p* |
| (Intercept) | 1.98 | 1.93 – 2.02 | <0.001 | 2.78 | 2.66 – 2.91 | <0.001 |
| General response [Strong] | 0.02 | -0.03 – 0.08 | 0.457 | 0.08 | -0.09 – 0.24 | 0.356 |
| General response [Weak] | 0.02 | -0.04 – 0.07 | 0.505 | -0.00 | -0.17 – 0.16 | 0.957 |
| Time since encounter [1st degree] | 3.41 | 0.55 – 6.26 | 0.019 | 48.38 | 43.99 – 52.77 | <0.001 |
| Time since encounter [2nd degree] | -0.90 | -3.34 – 1.53 | 0.466 | -7.98 | -10.55 – -5.40 | <0.001 |
| Previous encounter [Yes] | 0.07 | 0.00 – 0.13 | 0.038 | 0.12 | -0.06 – 0.31 | 0.194 |
| General response [Strong] × Time since encounter [1st degree] | 0.44 | -3.49 – 4.37 | 0.825 | 0.09 | -5.88 – 6.07 | 0.976 |
| General response [Weak] × Time since encounter [1st degree] | 2.63 | -1.22 – 6.48 | 0.180 | 1.84 | -4.11 – 7.78 | 0.545 |
| General response [Strong] × Time since encounter [2nd degree] | -1.65 | -5.01 – 1.72 | 0.338 | -2.55 | -6.05 – 0.96 | 0.154 |
| General response [Weak] × Time since encounter [2nd degree] | -2.23 | -5.50 – 1.04 | 0.182 | -2.39 | -5.89 – 1.10 | 0.180 |

Random Effects
|  |  |  |
| --- | --- | --- |
| $\sigma^2$ | 92.64 | 13.11 |
| N | 687<sub>EncounterID</sub> | 686<sub>EncounterID</sub> |
|  | 3<sub>Troop</sub> | 3<sub>Troop</sub> |
| Observations | 9524 | 9509 |
| Marginal R<sup>2</sup> | 0.000 | 0.020 |

**Table S7.**
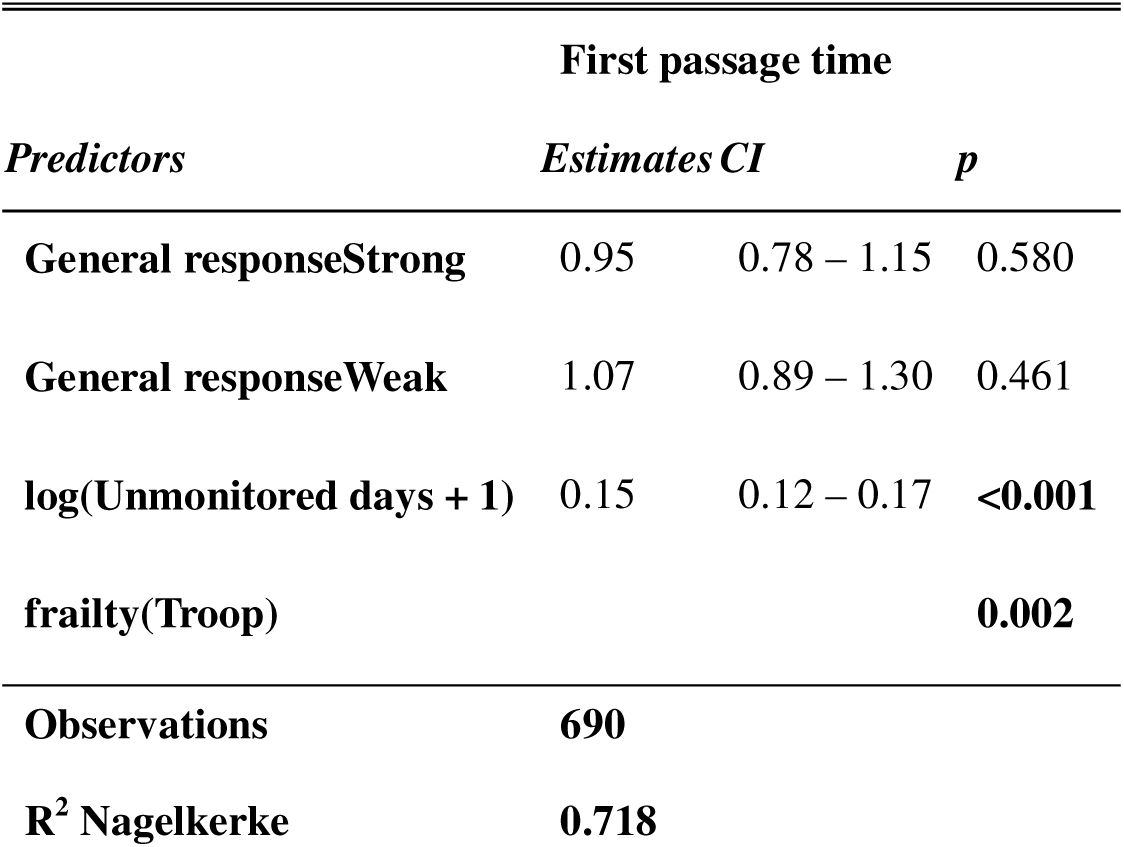
Output of the model 9 testing the time to return to a previous encounter (first passage time, in days) as a function of the troop’s general reaction level and during the encounter. CI = 95% confidence interval, *p* = p- value. frailty() indicates a random effect on the intercept of the predictor variable, included between the parentheses.

### Model Verifications

#### Generalised linear mixed models

We verified that the model assumptions were not violated by visually checking several distributions: the quantile-quantile distribution expected under the modelled distributions, the histogram of residuals, as well as the scatter plot of the fitted values *vs* the residuals using the *DHARMa* package (*92*). While the tests might be (too) sensitive, it visually indicated no major deviation from the required assumptions (Figure S1 to S7), which would threaten the validity of the results (*93*). When possible, we further assessed whether individual observations were disproportionately influencing our results using the “check_outliers” function of the *performance* package (*63*). No such influence was detected.

**Figure S1.**
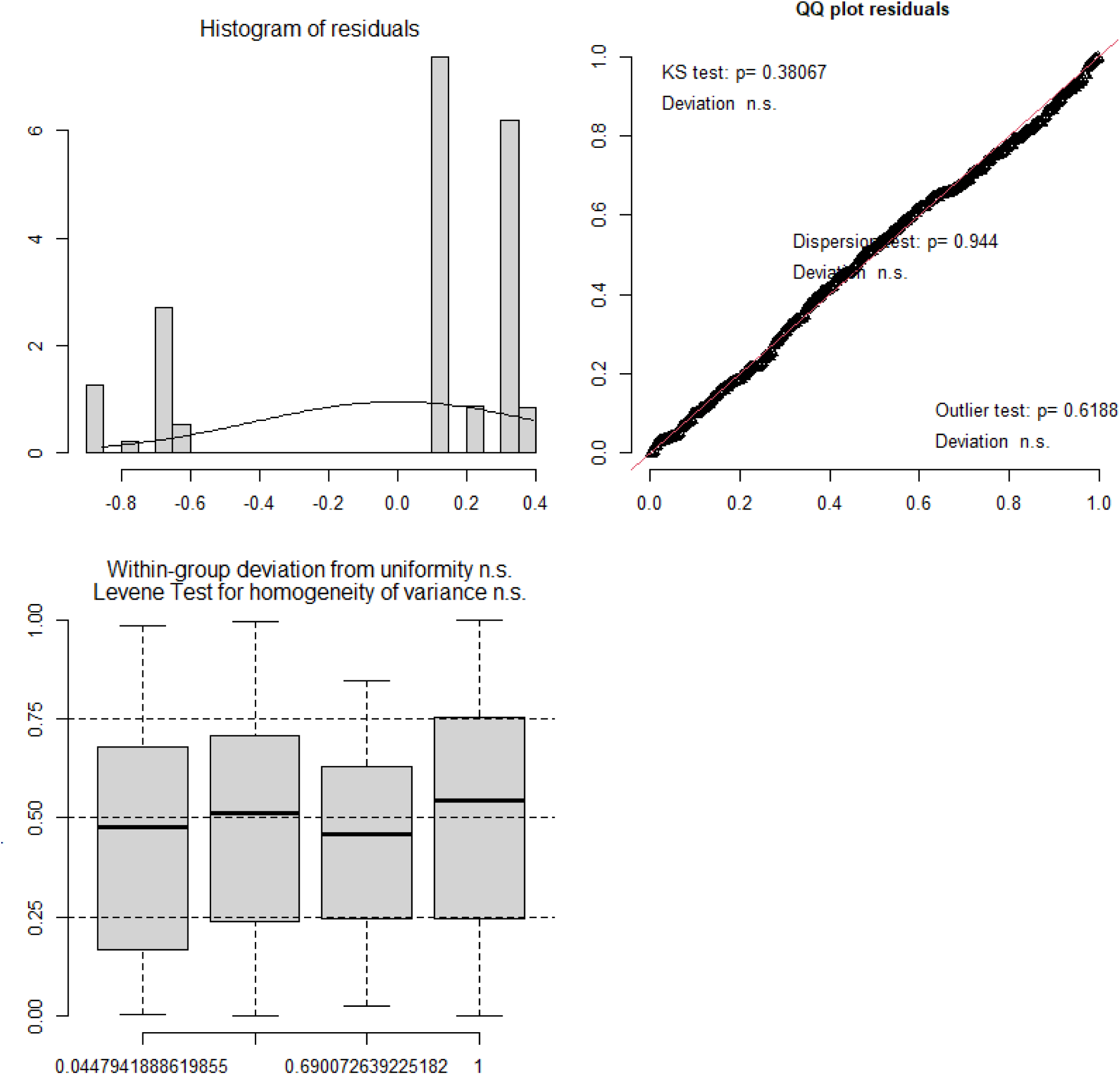
Model 1 (Alarm calling) assumptions check. Depicted are the histogram of residuals, the Q-Q plot (adapted for the modelled distribution; with test for deviation of the distribution (Kolmogorov-Smirnov, KS, outliers and overdispersion) and the scatter plot of the fitted values *vs* the residuals.

**Figure S2.**
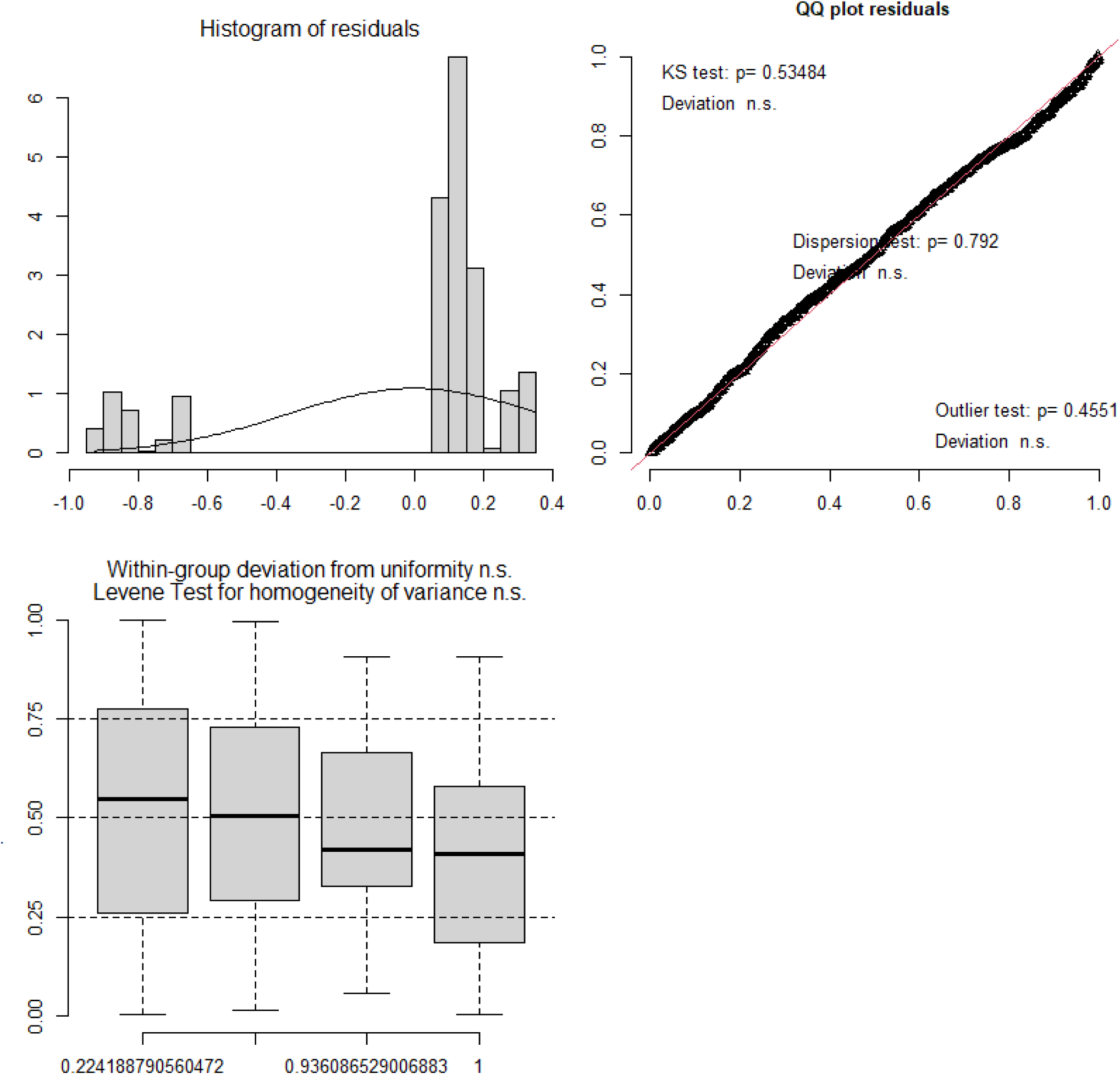
Model 2 (Alertness) assumptions check. Depicted are the histogram of residuals, the Q-Q plot (adapted for the modelled distribution; with test for deviation of the distribution (Kolmogorov-Smirnov, KS, outliers and overdispersion) and the scatter plot of the fitted values *vs* the residuals.

**Figure S3.**
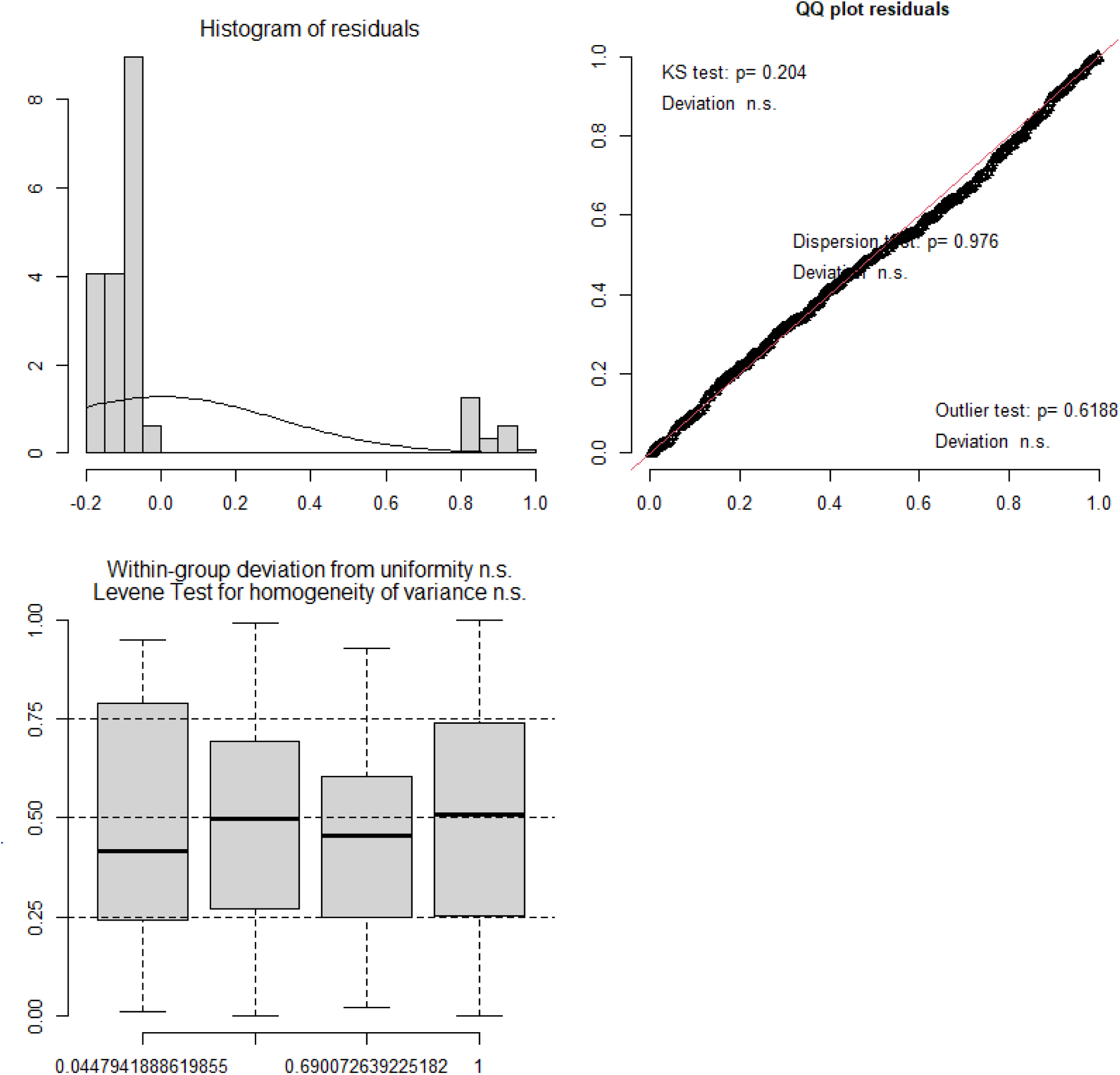
Model 3 (Agonistic reaction) assumptions check. Depicted are the histogram of residuals, the Q-Q plot (adapted for the modelled distribution; with test for deviation of the distribution (Kolmogorov-Smirnov, KS, outliers and overdispersion) and the scatter plot of the fitted values *vs* the residuals.

**Figure S4.**
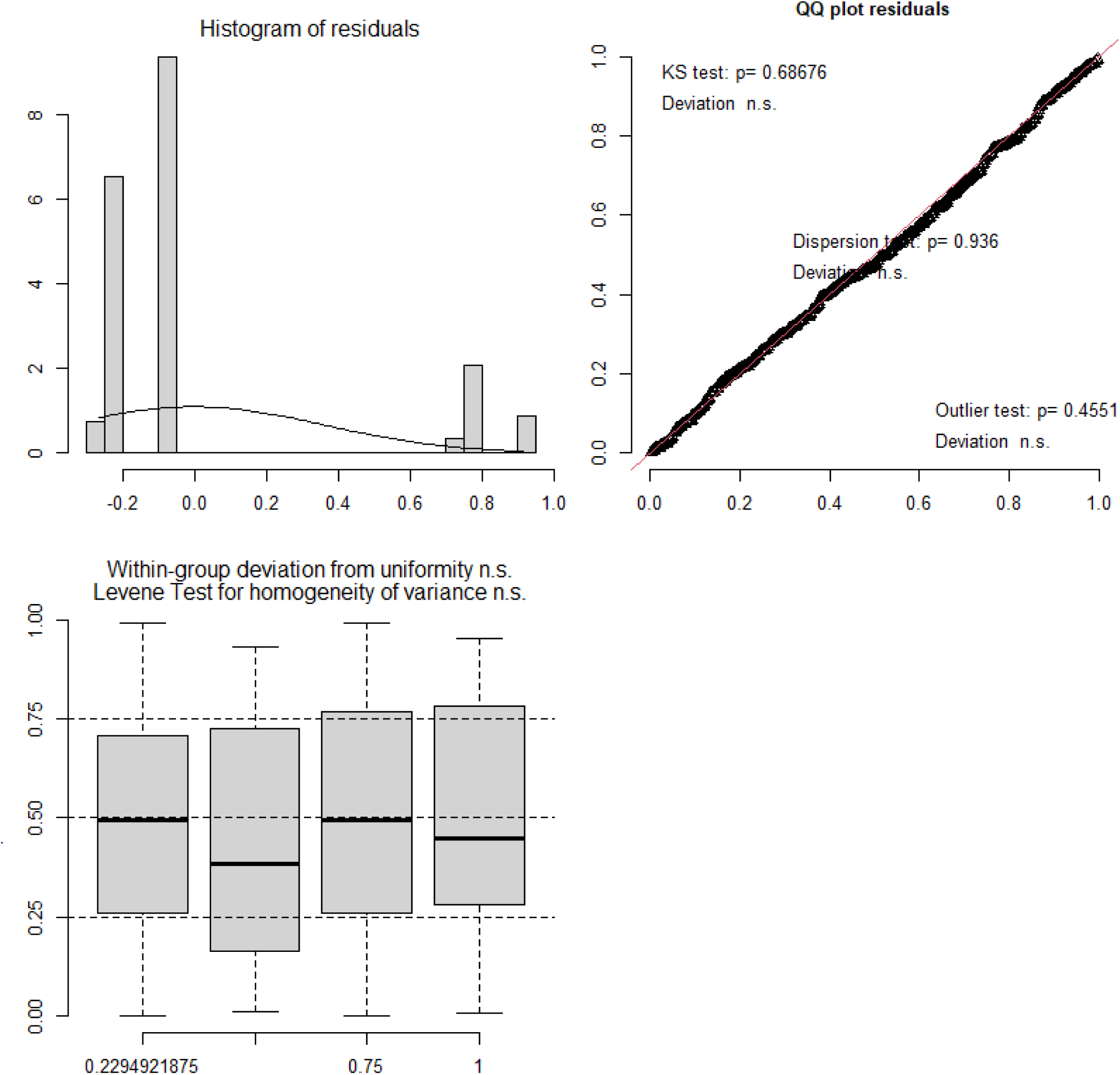
Model 4 (Fleeing) assumptions check. Depicted are the histogram of residuals, the Q-Q plot (adapted for the modelled distribution; with test for deviation of the distribution (Kolmogorov-Smirnov, KS, outliers and overdispersion) and the scatter plot of the fitted values *vs* the residuals.

**Figure S5.**
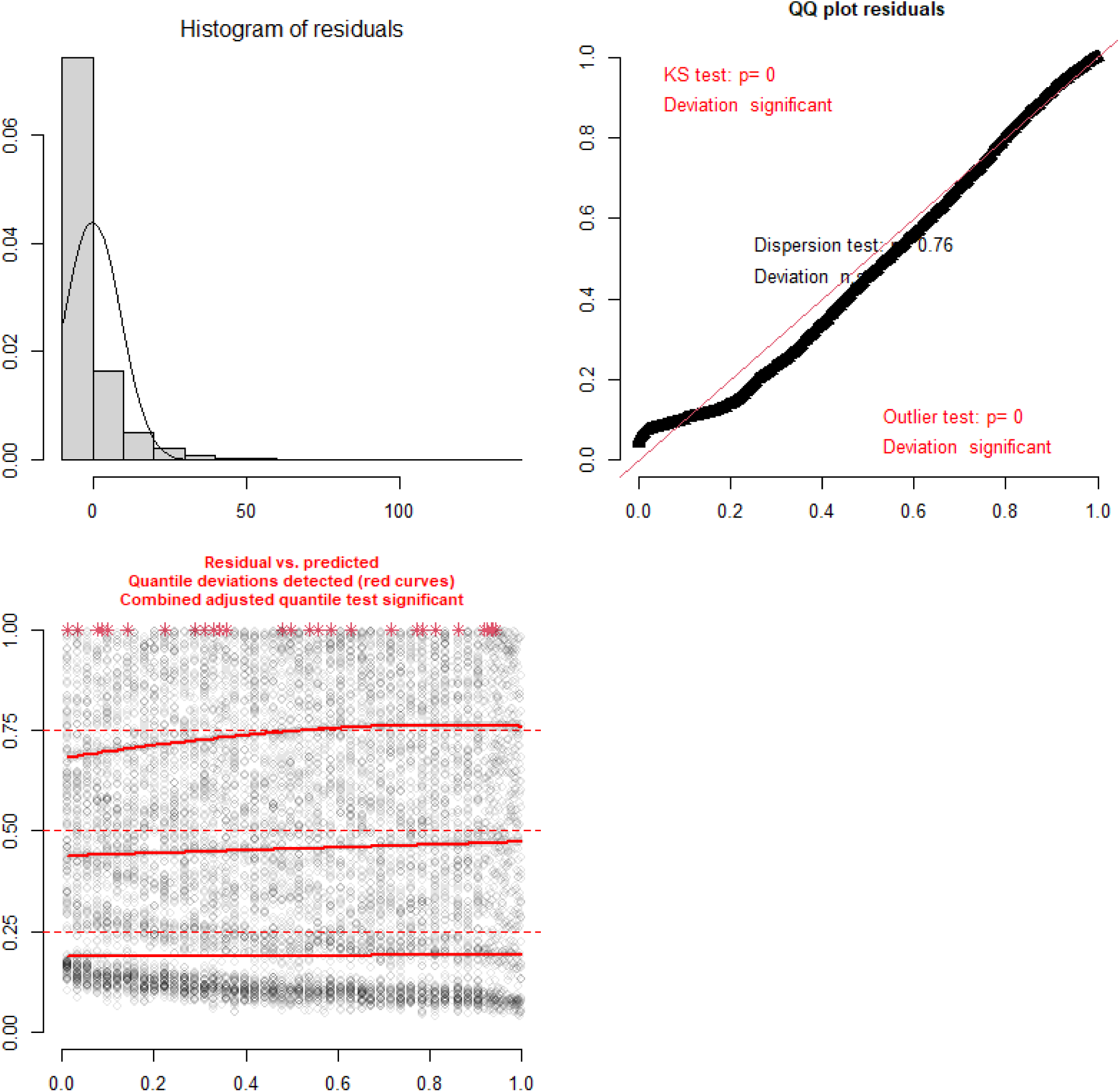
Model 6 (Movement speed) assumptions check. Depicted are the histogram of residuals, the Q-Q plot (adapted for the modelled distribution; with test for deviation of the distribution (Kolmogorov-Smirnov, KS, outliers and overdispersion) and the scatter plot of the fitted values *vs* the residuals.

**Figure S6.**
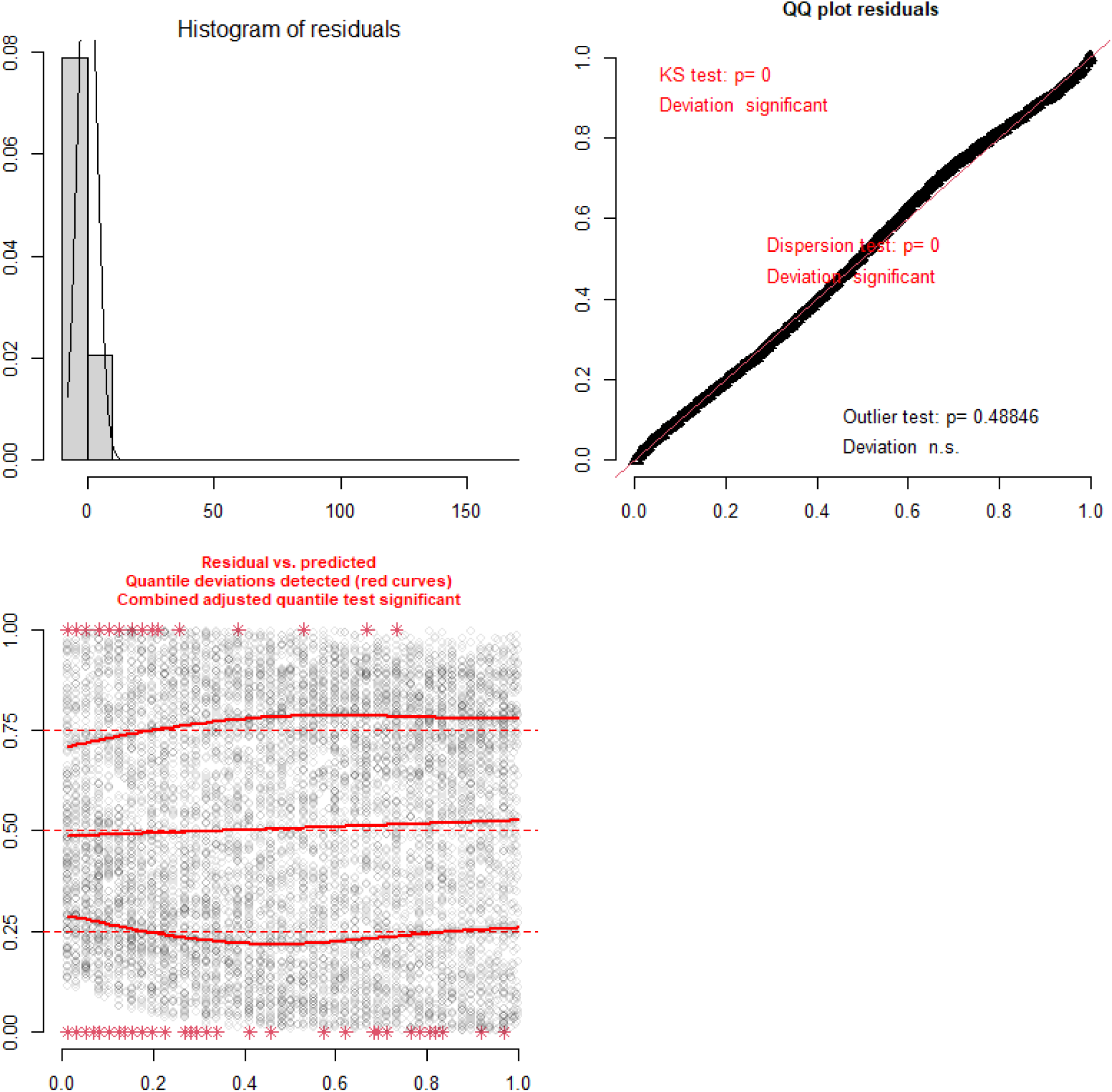
Model 7 (Displacement) assumptions check. Depicted are the histogram of residuals, the Q-Q plot (adapted for the modelled distribution; with test for deviation of the distribution (Kolmogorov-Smirnov, KS, outliers and overdispersion) and the scatter plot of the fitted values *vs* the residuals.

#### Proportional hazard models

We verified the proportional hazard hypothesis with the “cox.zph” function of the *survival* package (*61*, *62*), considering no random effect: this indicated no violation of the latter. We further verified the presence of outliers and influential points by checking the residuals of the models *vs* the fitted values, and the consequences on model estimations when observations were withdrawn one at a time (i.e., estimating the DfBetas, “resids” function, Figure S9). These analyses revealed no evidence of problematic outliers or influential points, and overall model fit was good, with a C-index above 0.7.

**Figure S7.**
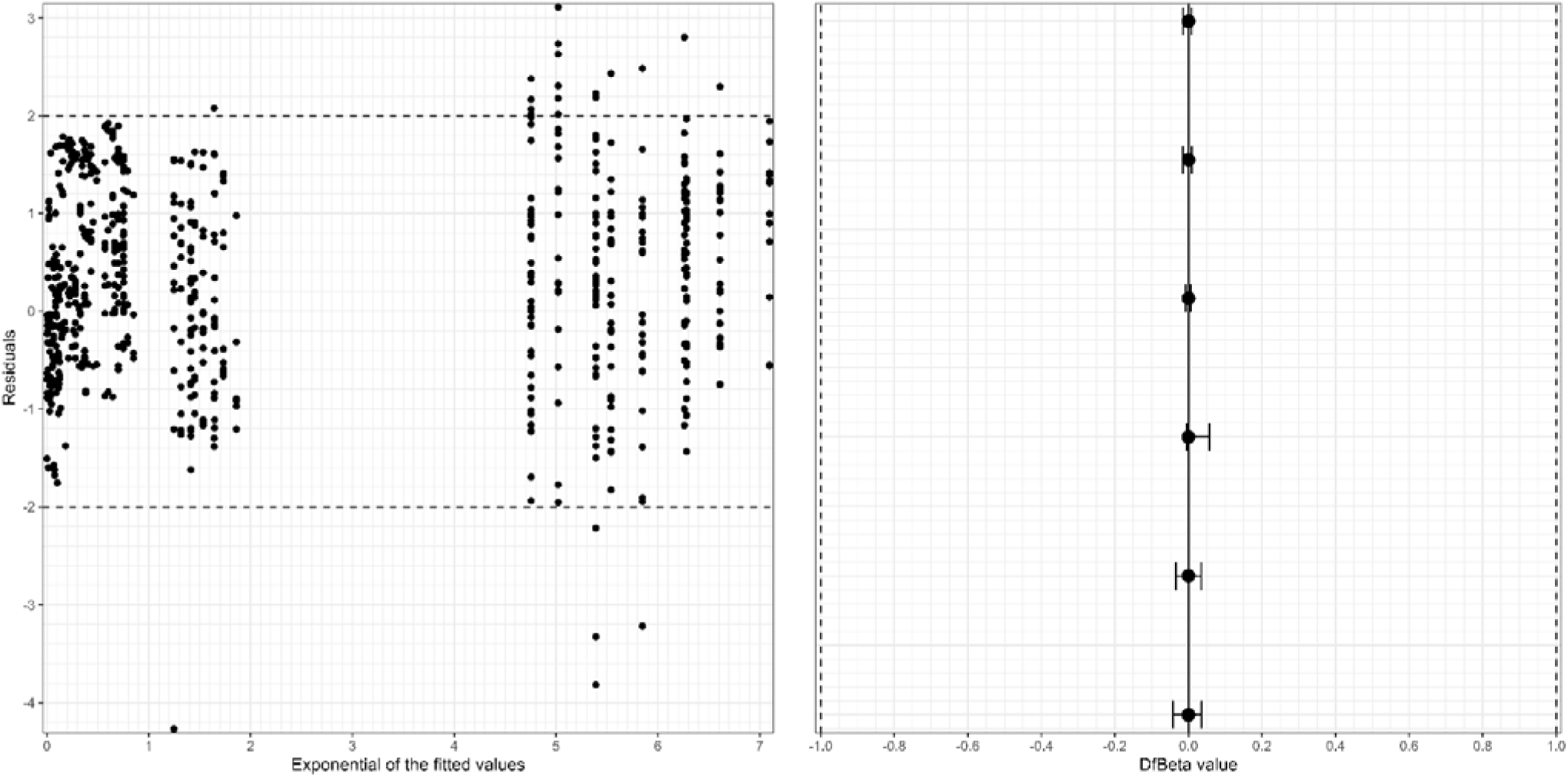
Visual check of the proportional hazard model fit. Left: Scatter plot of the residuals *vs* the fitted values. The dashed lines represent the upper and lower borders for which observations may be considered as outliers. Right: Visual representation of the DfBetas. Each circle and associated line correspond to the mean change and associated min-max range in the estimations when variables are removed one at a time, for each estimate of the model. The dashed lines represent the upper and lower borders for which changes may be considered substantial.

### Statistical artefact for the variable “dog number”?

Since the hierarchical rule creates a positive relationship between a dog’s risky label probability and dog number, we recomputed Model 5, focusing on the troop’s general response level as a function of the dog and context features. This time, rather than using a hierarchical rule, we set the label to the most common among dogs (which was possible for the variables “Leash,” “Healthy,” and “Size”). This rule should maintain an equal riskiest label probability over dog number, though the variance around this mean probability should decrease with dog number. Overall, the variable “Number” of dogs was not identified as a primary driver (likelihood ratio test: χ²L = 5.117, p = 0.08), and our conclusions were largely consistent (see Table S7 and Figure S8).

**Table S7.** Output of the model 6 investigating the drivers of the troop general response level using the majority rule (instead of the hierarchical rule).

| Level | Term | Estimate | Std.error | Statistic | p-value |
| --- | --- | --- | --- | --- | --- |
| None\|Weak | (Intercept) | -0.325 | 0.479 | -0.680 | 0.497 |
| Weak\|Strong | (Intercept) | **1.948** | **0.425** | **4.584** | **0.000** |
| None\|Weak | Number [>1] | **-0.502** | **0.233** | **-2.152** | **0.031** |
| Weak\|Strong | Number [>1] | -0.274 | 0.196 | -1.394 | 0.163 |
| None\|Weak | Size [Large] | **-1.227** | **0.239** | **-5.136** | **0.000** |
| Weak\|Strong | Size [Large] | **-0.594** | **0.216** | **-2.750** | **0.006** |
| None\|Weak | Healthy [Yes] | -0.033 | 0.292 | -0.113 | 0.910 |
| Weak\|Strong | Healthy [Yes] | -0.449 | 0.255 | -1.763 | 0.078 |
| None\|Weak | Chasing [Yes] | **-0.944** | **0.371** | **-2.542** | **0.011** |
| Weak\|Strong | Chasing [Yes] | **-0.979** | **0.288** | **-3.396** | **0.001** |
| None\|Weak | Barking [Yes] | -0.078 | 0.336 | -0.233 | 0.816 |
| Weak\|Strong | Barking [Yes] | -0.126 | 0.276 | -0.457 | 0.647 |
| None\|Weak | Location [Open] | **-1.832** | **0.402** | **-4.556** | **0.000** |
| Weak\|Strong | Location [Open] | **-0.674** | **0.300** | **-2.245** | **0.025** |
| None\|Weak | Leash [Yes] | 0.503 | 0.365 | 1.379 | 0.168 |
| Weak\|Strong | Leash [Yes] | 0.303 | 0.257 | 1.177 | 0.239 |
| None\|Weak | Distance<br>[Unreachable] | **1.690** | **0.301** | **5.619** | **0.000** |
| Weak\|Strong | Distance<br>[Unreachable] | 0.182 | 0.204 | 0.894 | 0.371 |
|  | PreviousEncounter<br>[Yes] | -0.300 | 0.250 | -1.199 | 0.231 |

**Figure S8.**
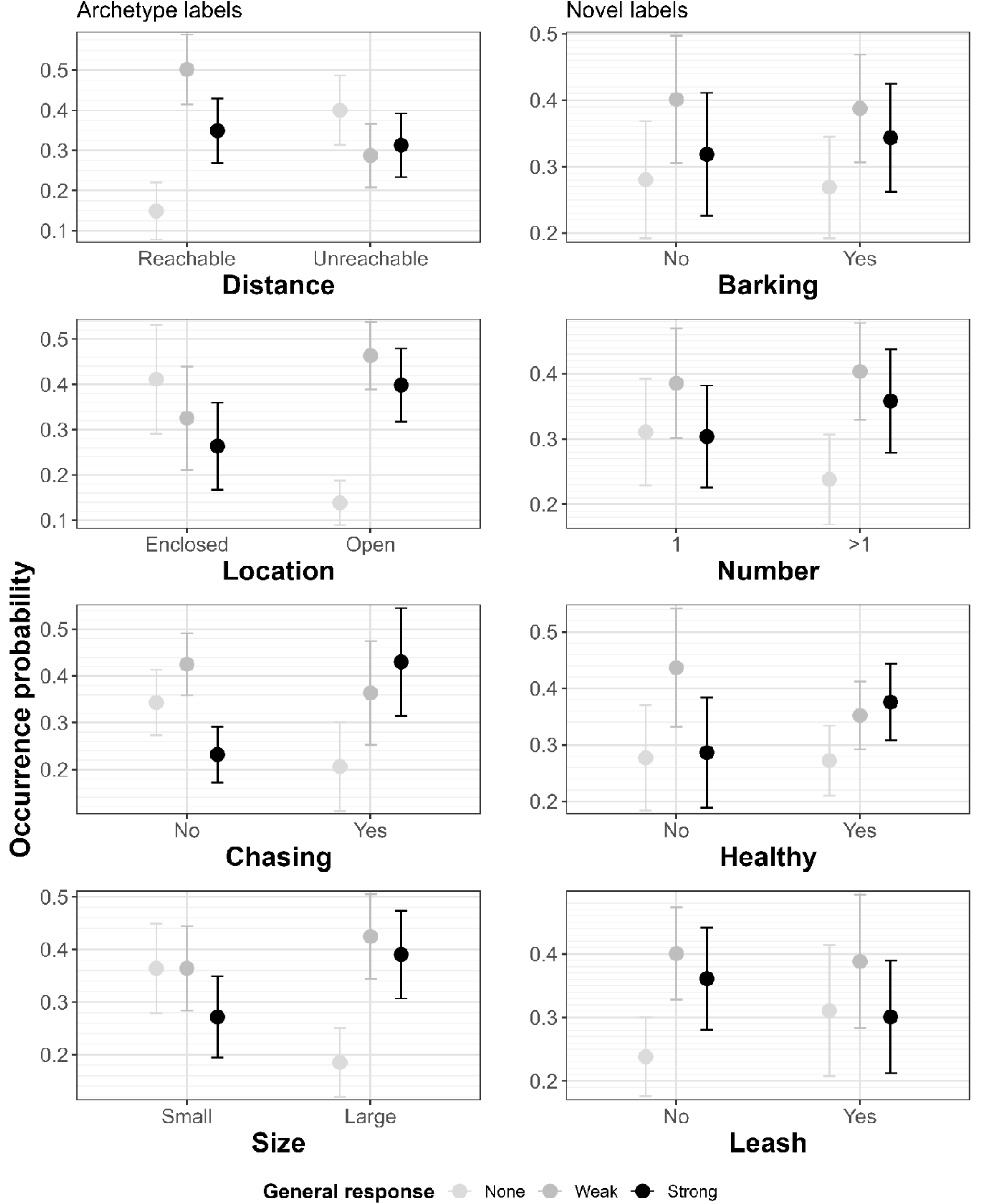
Troop general response level as a function of dogs’ features and encounter micro-habitat context using a majority rule instead of a hierarchical rule. The plain circles represent the model marginal estimates, while the segments represent the associated 95% confidence intervals.

